# Early adipose tissue wasting in a novel preclinical model of human lung cancer cachexia

**DOI:** 10.1101/2024.09.27.615385

**Authors:** Deena B. Snoke, Jos L. van der Velden, Emma R. Bellafleur, Jacob S. Dearborn, Sean M. Lenahan, Alexandra E. Beal, Reem Aboushousha, Skyler C. J. Heininger, Jennifer L. Ather, Madeleine M. Mank, Hailey Sarausky, Daniel Stephenson, Julie A. Reisz, Angelo D’Alessandro, Devdoot Majumdar, Thomas P. Ahern, Kaiwen Xu, Kim L. Sandler, Bennett A. Landman, Yvonne M.W. Janssen-Heininger, Matthew E. Poynter, David J. Seward, Michael J. Toth

**Affiliations:** Department of Medicine, University of Vermont College of Medicine, Burlington, VT; Department of Pathology and Laboratory Medicine, University of Vermont College of Medicine, Burlington, VT; Department of Surgery, University of Vermont College of Medicine, Burlington, VT; University of Vermont Cancer Center, University of Vermont College of Medicine, Burlington, VT; Department of Biochemistry and Molecular Genetics, University of Colorado Anschutz Medical Campus, Aurora, CO; Department of Computer Science, Vanderbilt University and Vanderbilt University Medical Center, Nashville, TN, USA; Department of Biomedical Engineering, Vanderbilt University and Vanderbilt University Medical Center, Nashville, TN, USA; Department of Radiology, Vanderbilt University and Vanderbilt University Medical Center, Nashville, TN, USA

**Keywords:** Lung neoplasms, lipolysis, skeletal muscle, organoids, inflammation

## Abstract

Cancer cachexia (CC), a syndrome of skeletal muscle and adipose wasting, reduces responsiveness to therapies and increases mortality. There are no approved treatments for CC, which may relate to discordance between pre-clinical models and human CC. To address the need for clinically relevant models of lung CC, we generated inducible, lung epithelial cell specific *Kras^G12D/+^* (*G12D*) mice. *G12D* mice develop CC over a protracted time course and phenocopy tissue and tumor, cellular, mutational, transcriptomic, and metabolic characteristics of human lung CC. *G12D* mice demonstrate early loss of adipose, a phenotype that was apparent across numerous models of CC and translates to patients with lung cancer. Tumor-released factors promote adipocyte lipolysis, a driver of adipose wasting in CC, and adipose wasting was inversely related to tumor burden. Thus, *G12D* mice model key features of human lung CC and highlight a role for early tumor metabolic reprogramming of adipose tissue in CC.

**Graphical Abstract:** 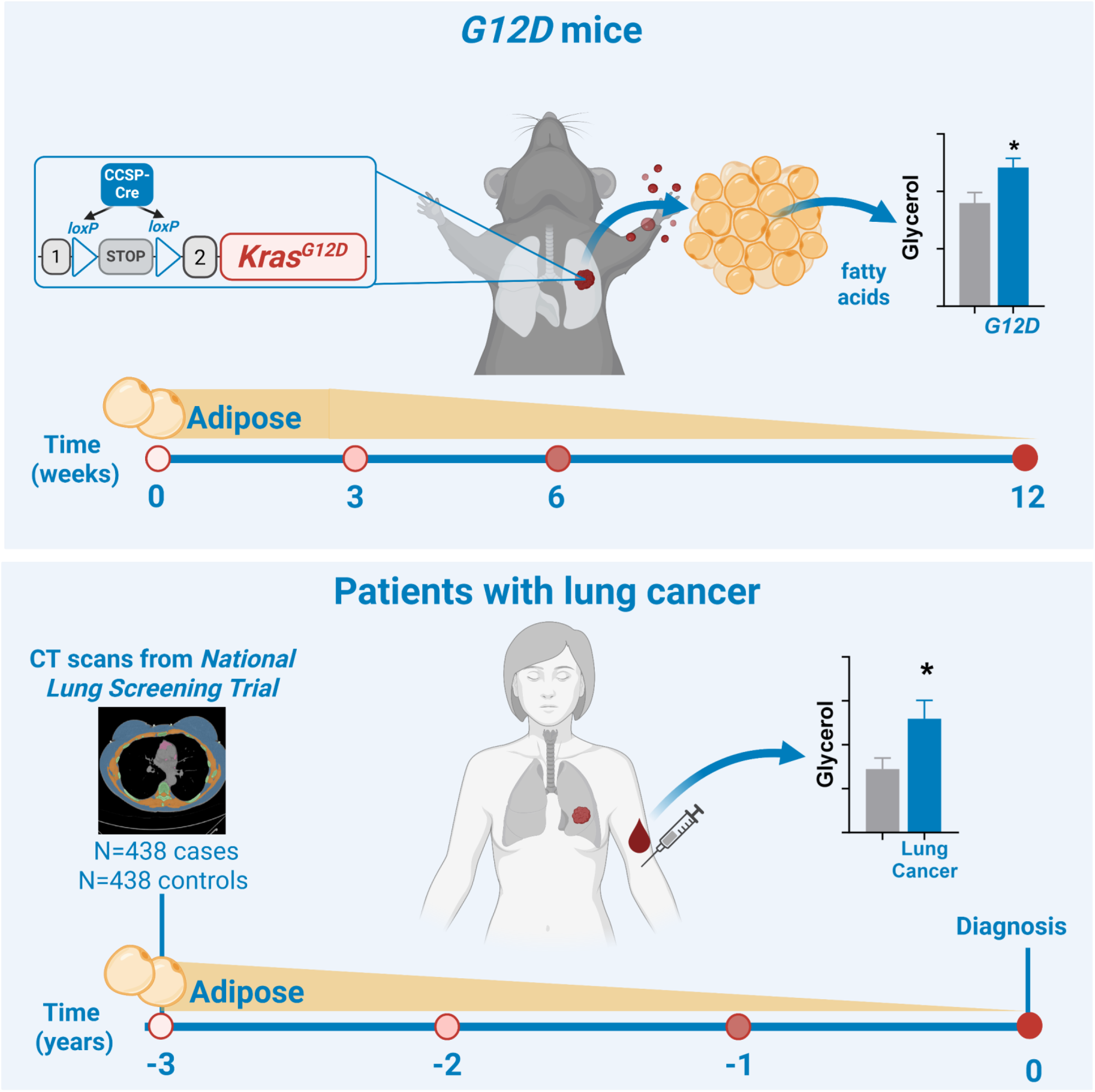

## Introduction

Cancer cachexia (CC) is characterized by unintentional loss of body weight ^1^ and contributes to poor response to treatments and related toxicities and increased morbidity and mortality ^1–11^. Animal models have served as the basis for knowledge of the pathophysiology of CC ^12^ because the syndrome typically occurs in aggressive cancers and later-stage disease, where patients are unwilling or unable to participate in research. While these models have generated a wealth of data ^12^, there are no effective, approved treatments for CC. This lack of translating evidence into treatments may be explained, in part, by the fact that current models do not mimic human CC ^13–21^.

Lung cancer is the second most common cancer and the leading cause of cancer-related death ^22, 23^. Most patients with lung cancer exhibit signs of CC ^24, 25^, and CC reduces mean survival time and responsiveness to chemo- and immunotherapies ^11, 26^. Most preclinical data on lung CC are derived from the murine Lewis lung carcinoma (LLC) model, in which tumor cells are injected subcutaneously and grow rapidly. Weight loss begins when the tumor reaches ∼5% of body weight, and mice progress to >10% body weight loss in 1-3 weeks ^27^. Neither tumor growth, location, or size, nor weight loss characteristics of this model reflect lung CC in humans ^28^. Thus, the field would benefit from new models of lung CC that better emulate human lung CC.

To address this need, we developed mice with an activatable lung epithelial cell-specific *Kras^G12D^* allele (*G12D*) to evaluate its utility as a model of lung CC. We report that *G12D* mice display classic adipose and skeletal muscle tissue wasting phenotypes of CC over a protracted time course (∼12 weeks) and phenocopy tumor and tissue cellular, mutational, transcriptomic, and metabolic characteristics relevant to human lung CC. Additionally, adipose tissue wasting drove CC initiation in both singular *Kras^G12D^* mutation and when combined with other driver mutations (*e.g., Stk11^-/-^*), was inversely related to tumor burden, and could be incited by tumor released factors. In a large, prospective cohort study, we confirmed the clinical relevance of early adipose tissue wasting. Thus, early tumor metabolic reprogramming of adipose tissue may be important for CC initiation and tumor biology.

## Results

### Lung epithelial cell (club cell)-specific Kras^G12D^ induction causes CC

Induction of the *Kras^G12D^* allele with tamoxifen administration (**Fig. 1A**) promotes synchronous pulmonary lesions originating in the terminal bronchioles, the predominant anatomic location of club cells (**Fig. 1B**). Lesions exist along the histological spectrum of epithelial proliferation, ranging from florid bronchial hyperplasia to adenoma to atypical adenomatous hyperplasia to adenocarcinoma, with no evidence for dissemination to major organs or lymph nodes. Lung adenocarcinomas were confirmed histologically at the initiation of weight loss (6 weeks post-induction; Fig. 1C) within the spectrum of synchronous pulmonary lesions characterized by marked cellular atypia, architecturally distorted glands, the absence of cilia, and enhanced mitotic index (**Fig. S1A**). In fact, the histologic hallmarks of adenocarcinoma were present, but not as pronounced, as early as 3.5 weeks post-induction (**Fig. S1B**). Next, we examined whether CCSP promoter activity was evident only upon injection of tamoxifen, as non-tamoxifen-inducible (*i.e.,* leaky) Cre expression from the CCSP promoter has been observed ^29^. In line with these reports, in *G12D* mice not receiving tamoxifen, we found histological evidence for lung lesion development (**Fig. S2**), albeit much less severe than in mice with tamoxifen injection. Because of this, we used wild-type, *Scgb1a1-CreER^TM^* littermates (WT; *Kras^+/+^*) injected with tamoxifen as controls, which has the advantage of controlling for toxicity and off-target effects of intracellular Cre activation and tamoxifen, respectively. Finally, tumor growth can affect the function of the organ where it resides, promoting weight loss independent of the tumor. In *G12D* mice, while animals demonstrate labored breathing and reduced activity level at advanced stage CC (weeks 10-12), they show no evidence of labored breathing or reduced activity at the initiation of body weight loss at 6 weeks (Fig. 1C). To evaluate whether there is evidence for impaired lung function that may drive CC, we measured blood oxygen saturation at 3 weeks post-induction, when body weight begins to diverge between groups (Fig. 1C). We found no evidence for hypoxemia (**Fig. S3**), suggesting that respiratory dysfunction in *G12D* mice is unlikely to contribute to CC.

**Figure 1.**
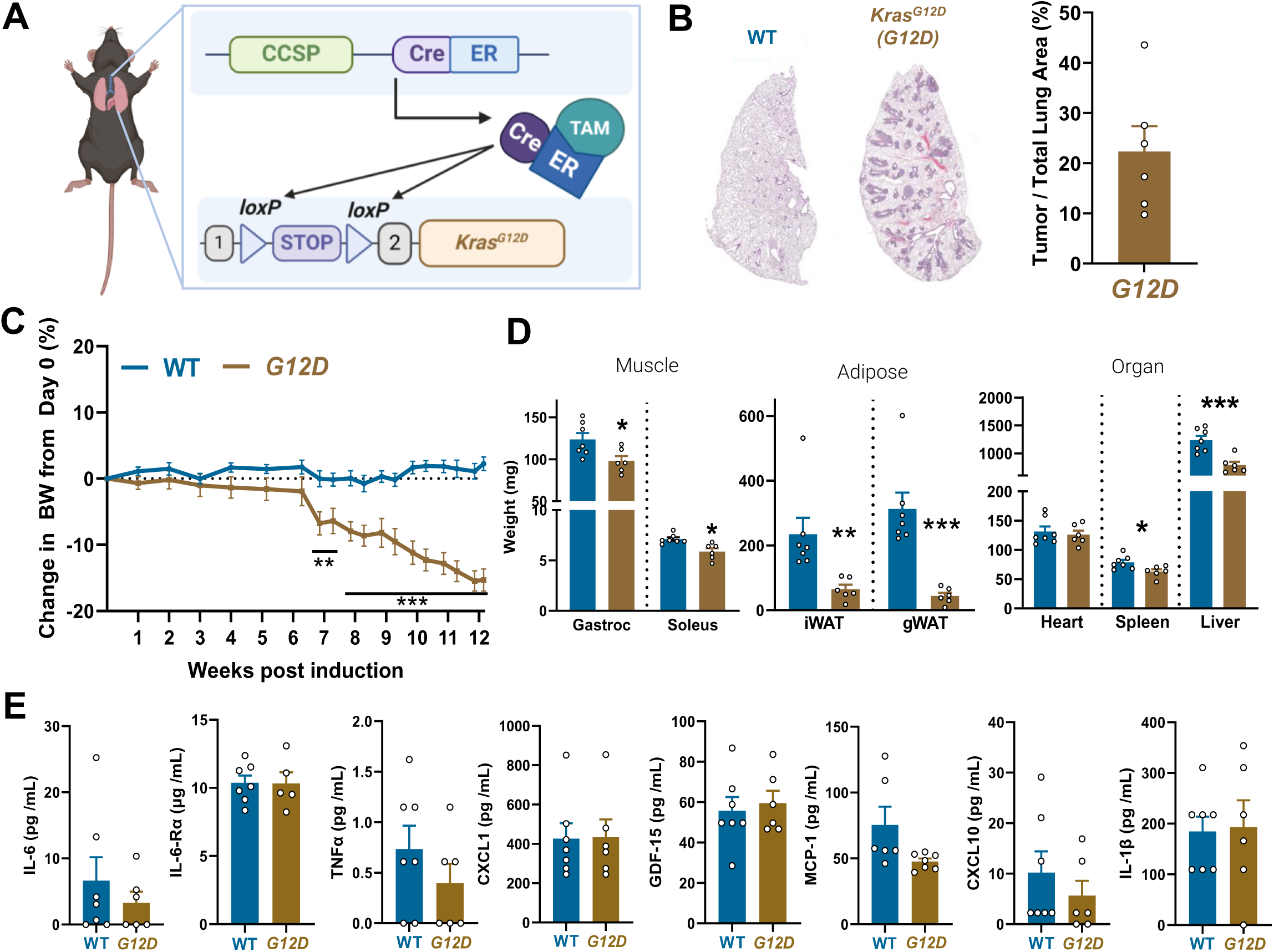
Characterization of lung epithelial cell specific, inducible Kras^G12D/+^ (G12D) mice at 12 weeks post-induction. *Kras^G12D/+^ (G12D)* mice harbor a tamoxifen (TAM)-inducible lung epithelial cell specific *Kras^G12D^* mutation under the club cell secretory protein (CCSP) promoter (A). Four-to 5-month-old mice received intraperitoneal TAM, leading to adenocarcinoma development in lung bronchioles by 12 weeks post-induction compared with wild-type (WT) mice and average tumor burden (percent tumor area / total lung area) (B). Change in body weight relative to Day 0 (C), skeletal muscle, fat pad and organ weights (D) and serum inflammatory cytokines (interleukin 6 (IL-6), soluble IL-6 receptor alpha (IL-6Rα), tumor necrosis factor alpha (TNFα), CXC motif ligand 1 (CXCL1/KC), growth differentiation factor 15 (GDF-15), monocyte chemoattractant protein 1/C-C motif ligand 2 (MCP-1/CCL2), CXCL10, and interleukin-1 beta (IL-1β)) in wild-type (WT; n=7; blue bars/lines) and G12D mice (n=6; gold bars/lines)(E). Data were analyzed using repeated measures ANOVA (C) or Student’s t-test (D, E), with * p<0.05, ** p<0.01, and *** p< 0.001. Note that n=5-7/group for serum cytokine measures. Image created with Biorender.com (A). See also: supplemental figures S1-4.

*G12D* mice began losing weight 6 weeks post-induction (**Fig. 1C**), independent of differences in food intake or fecal energy content (**Fig. S4A-B**). By 12 weeks, *G12D* mice experienced an average body weight loss of 15%, with most animals (83%) displaying CC (>10% body weight loss). Weight loss in *G12D* mice was explained by loss of skeletal muscle (gastrocnemius and soleus), white adipose tissue (inguinal (iWAT) and perigonadal (gWAT)), and organ (spleen and liver) weight (**Fig. 1D**). No group differences were found in serum inflammatory cytokine concentrations (**Fig. 1E**).

### Skeletal muscle and adipose tissue cellular wasting in G12D mice

To further characterize tissue wasting, myofiber size was assessed at 12 weeks post-induction. *G12D* mice exhibited reduced gastrocnemius (**Fig. 2A, Fig. S5A**), but not soleus, muscle fiber cross-sectional area (CSA; **Fig. 2A**, **Fig. S5B**). Gastrocnemius muscle of *G12D* mice showed greater expression of key genes involved in protein breakdown that are implicated in CC (*Fbxo32, Becn1*), and *Il6, Tnf,* and *Lif* mRNA compared with WT (**Fig. 2B**), with cytokines indicating local inflammation that may drive muscle wasting^30^. In soleus, mRNA levels of *Il6* were greater in *G12D* mice, with no differences in proteolytic or inflammation markers (**Fig. S5C**).

**Figure 2.**
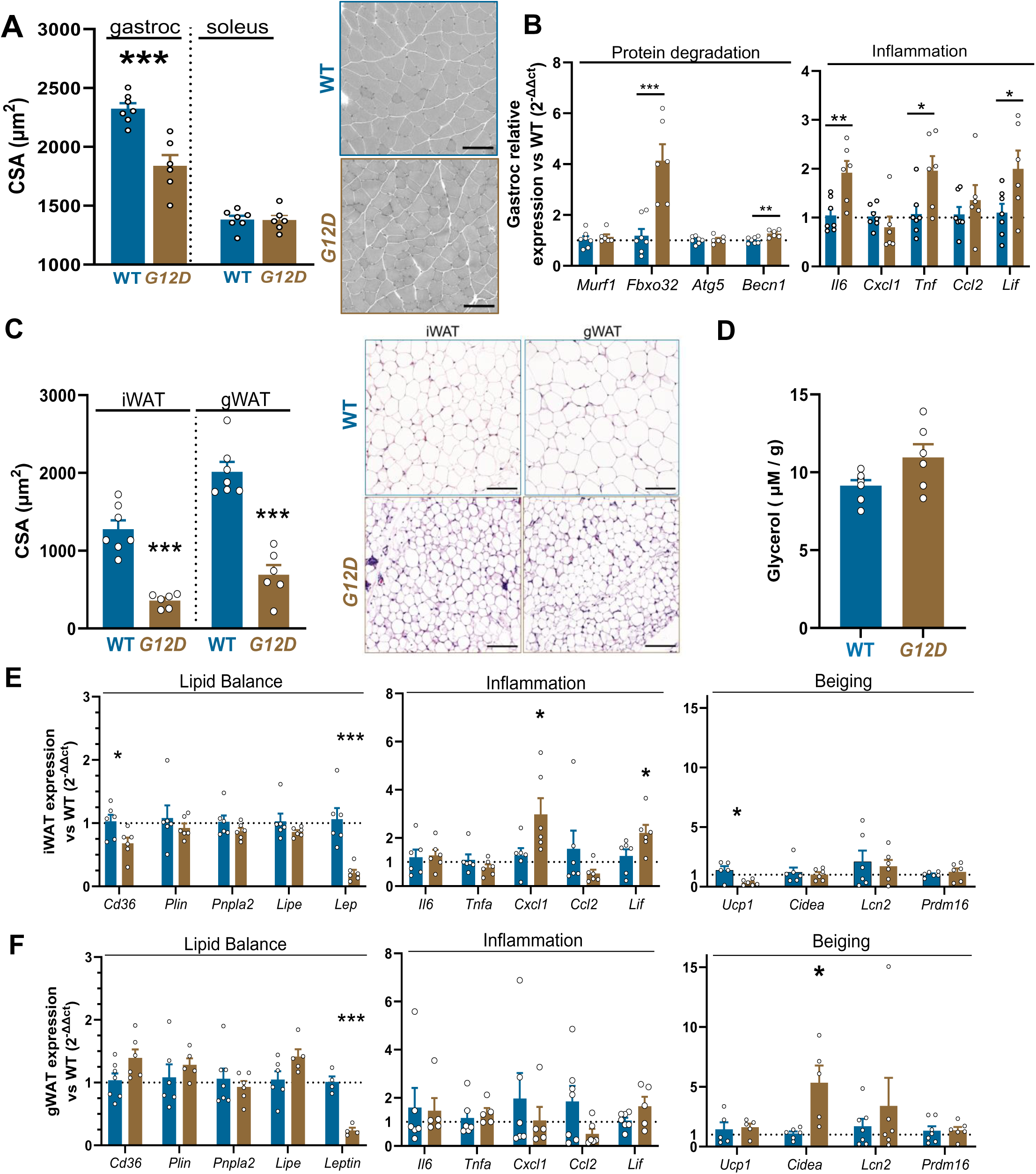
Effect of lung epithelial cell specific, inducible Kras^G12D/+^ (G12D) on skeletal muscle and adipose tissue wasting and mRNA expression at 12-weeks post-induction. Gastrocnemius (gastroc) and soleus fiber cross-sectional area (CSA) 12 weeks post-induction in wild-type (WT; n=7; blue bars) and *Kras^G12D/+^* (*G12D*) mice (n=6; gold bars) and representative images (scale bars=100 μm) (A). mRNA levels for markers of protein degradation and inflammatory cytokines in gastrocnemius muscle, expressed relative to the WT (B). Cross-sectional area (CSA) of adipocytes from inguinal (iWAT) and gonadal white adipose tissue (gWAT) depots in WT (n=7) and *G12D* (n=6) mice and representative sections (scale bars= 100 μm) (C). Serum glycerol levels in WT and *G12D* mice standardized to body mass (D). Gene expression of iWAT (E) and gWAT (F) depots for markers of lipid balance, inflammation, and beiging. Data were analyzed by Student’s t-test, with * *p*<0.05 and *** *p*<0.001. Significance values where *p*>0.05. See also: supplemental figures S5-7.

Subcutaneous iWAT and visceral gWAT adipocytes exhibited markedly lower CSA at 12 weeks post-induction in *G12D* mice compared with WT mice (**Fig. 2C**), and we observed higher circulating medium and long-chain fatty acids and markers of carnitine/fatty acid metabolism (**Fig. S6**), as well as a trend (*p*=0.06) toward greater circulating glycerol levels (**Fig. 2D**). In *G12D* iWAT, mRNA of the lipid transporter, *Cd36,* and mitochondrial uncoupling marker, *Ucp1,* were lower, and cytokines, *Cxcl1* and *Lif,* greater compared with WT (**Fig. 2E**). In gWAT of *G12D* mice, we observed greater expression of mRNA for the beiging marker, *Cidea*, and trends toward greater *Cd36* (*p*=0.06) and *Lipe* (*p*=0.07; **Fig. 2F**), the latter being the transcript of hormone-sensitive lipase. As expected, mRNA levels of *Lep* (leptin) were lower in iWAT and gWAT of *G12D* mice compared with WT mice (**Fig. 2E-F**), reflecting reduced adipocyte size. Of note, controlling for sex did not alter group differences in adipose and muscle cell size and tissue weights. Collectively, these data show that *G12D* mice exhibit tissue, cellular, transcriptomic, and metabolomic hallmarks of CC muscle and adipose wasting.

Similar driver mutations can elicit different tumor metabolic phenotypes in epithelial cells in different organs (lung vs. pancreas ^31^), which, in turn, may affect the nature of CC. We chose to express *Kras^G12D^* in club cells because this is a recognized model of lung adenocarcinoma^32^, but adenocarcinoma can also develop in alveolar type-2 (AT2) cells^33^. Thus, we evaluated mice expressing *Kras^G12D^* in AT2 cells following induction via adenovirus that permits AT2-specific Cre recombinase expression (Ad5-SPC-Cre). At ∼12 weeks post-induction, we found no differences in skeletal muscle, adipose depot, or organ weights compared to animals administered control adenovirus (**Fig S7A**). We also activated the *Kras^G12D^* allele using an adenovirus that permits epithelial cell agnostic Cre recombinase expression (Ad5-CMV-Cre). In these mice, we found reduced weights of two adipose depots and the spleen and patterns towards reductions in other adipose depots, albeit non-significant, but no differences in skeletal muscle weights (**Fig S7A**), suggesting cell type-specific differences in the ability of *Kras^G12D^*to incite CC. The general absence of overt CC wasting phenotypes in mice with both IT Ad-Cre induction approaches likely related to reduced tumor burden, as we found relative tumor burdens in Ad-Cre mice equivalent to our *G12D* mice at 3 weeks (∼5%; **Fig S7B**). Thus, the induction method (*i.e.,* intraperitoneal tamoxifen vs. oropharyngeal adenovirus expression Cre recombinase) affects the timing of CC, although the phenotype of early adipose wasting appears to be similar when considered relative to tumor burden.

### Early tissue wasting phenotypes in G12D mice

Utilizing the protracted time course of wasting in *G12D* mice, we evaluated *G12D* mice at 6 weeks post-induction, at the initiation of weight loss (Fig. 1C), to examine tumor and peripheral tissue morphological, transcriptomic, and metabolic features of CC initiation. Tumor burden averaged 10.5 ± 1.1% (**Fig. 3A-B**) while *G12D* mice lost an average of 5% of their body weight (**Fig. 3C**). Assuming an average lung weight of 200 mg, we estimate tumor burdens of ∼0.08% and ∼0.16% of body weight at 6-(Fig. 3B) and 12 (Fig 1B) weeks post-induction, respectively, which is closer to human lung cancer than other lung CC models. No differences in skeletal muscle or organ weights were noted between groups 6 weeks post-induction, but *G12D* mice showed markedly lower iWAT and gWAT weights (**Fig. 3D**). In a separate group of animals, we assessed other skeletal muscles (plantaris, tibialis anterior, diaphragm, and extensor digitorum longus) and found no differences in their weights (**Fig S8A**). Finally, no differences were found in a broad range of circulating cytokines (**Fig. 3E; Fig S8B**).

**Figure 3.**
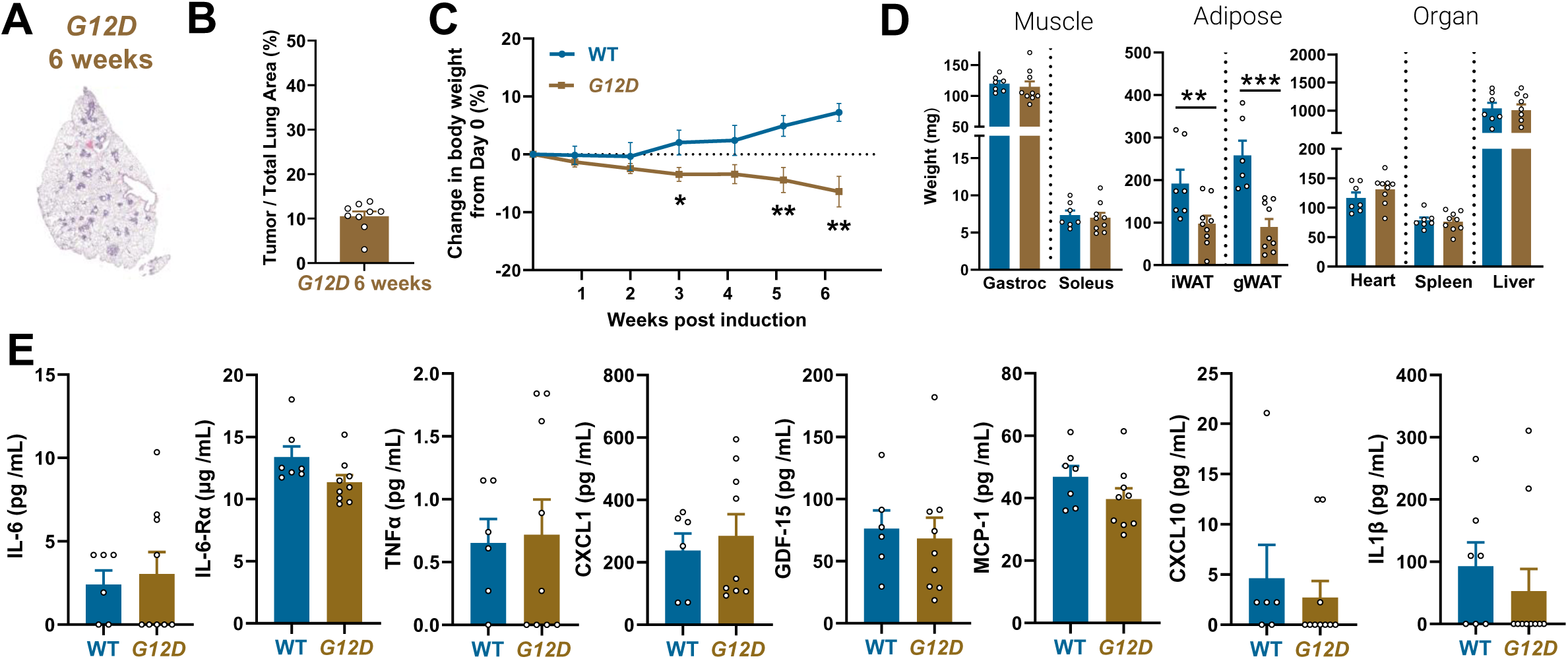
Characterization of lung epithelial cell specific, inducible Kras^G12D/+^ (G12D) mice CC at 6-weeks post-induction. Representative lung cross-section of a *Kras^G12D/+^* (*G12D*) mouse (A) and tumor burden (percent tumor area / total lung area) (B). Change in body weight relative to Day 0 in wild-type (WT; n=7; blue lines/bars) and *G12D* (n=8; gold lines/bars) mice (C). Muscle, fat pad, and organ weights of 6-week WT and *G12D* mice (D). Serum inflammatory cytokines in 6-week mice (E). Data were analyzed by repeated measures analysis of variance (C) and Student’s t-tests (D, E), with ** *p*<0.01 and *** *p*<0.001. See also: supplemental figure S8.

At the cellular level, consistent with skeletal muscle weights (Fig. 3D), neither gastrocnemius nor soleus muscle fiber CSA differed between groups at 6 weeks post-induction (**Fig. 4A**, **Fig. S8C-D**). Moreover, except for a trend (*p*=0.06) towards statistically significant greater *Fbxo32* mRNA in the gastrocnemius of *G12D* mice compared with WT mice, no differences in gene expression for markers of protein degradation or inflammatory cytokines were observed (**Fig. S7E-F**). Additionally, we found no group differences in tibialis anterior muscle fiber CSA (**Fig S8G**). In contrast, *G12D* mice showed marked atrophy of adipocytes in both depots at 6 weeks post-induction (**Fig. 4B**). In *G12D* mice, gene expression markers of lipid balance (*Plin1, Pnpla2,* and *Lipe*) were elevated in iWAT (**Fig. 4C**), with a similar expression pattern in *Plin1* and *Pnpla2* (both *p*=0.09) in gWAT compared with WT mice (**Fig. 4D**). In both depots, we found no differences in mRNA levels of inflammatory cytokines. However, as local tissue inflammation may contribute to CC ^30^, we conducted a more in-depth, unbiased assessment of inflammatory cytokine expression in gWAT at 3 and 6 weeks post-induction by bulk RNA-seq (Fig 4E). At 3 weeks post-induction in gWAT, *G12D* gWAT showed 20 differentially expressed genes (DEG; 7 upregulated and 13 downregulated) compared with WT gWAT (**Fig S9A**). Pathway enrichment analysis showed no evidence for upregulation of inflammatory pathways/signaling (**Fig S9B**). In fact, inflammatory response and cytokine signaling were downregulated. By 6 weeks post-induction in gWAT, *G12D* gWAT showed 31 differentially expressed genes (DEG; 17 upregulated and 14 downregulated) compared with WT gWAT (**Fig S9C**). Pathway analysis showed upregulation of inflammatory response and numerous cytokine signaling pathways (**Fig S9D**). Thus, adipose tissue inflammation develops coincident with CC-associated wasting, which suggest the possibility that, like muscle ^30^ (Fig 2B), adipose wasting may be modulated by local tissue inflammation.

**Figure 4.**
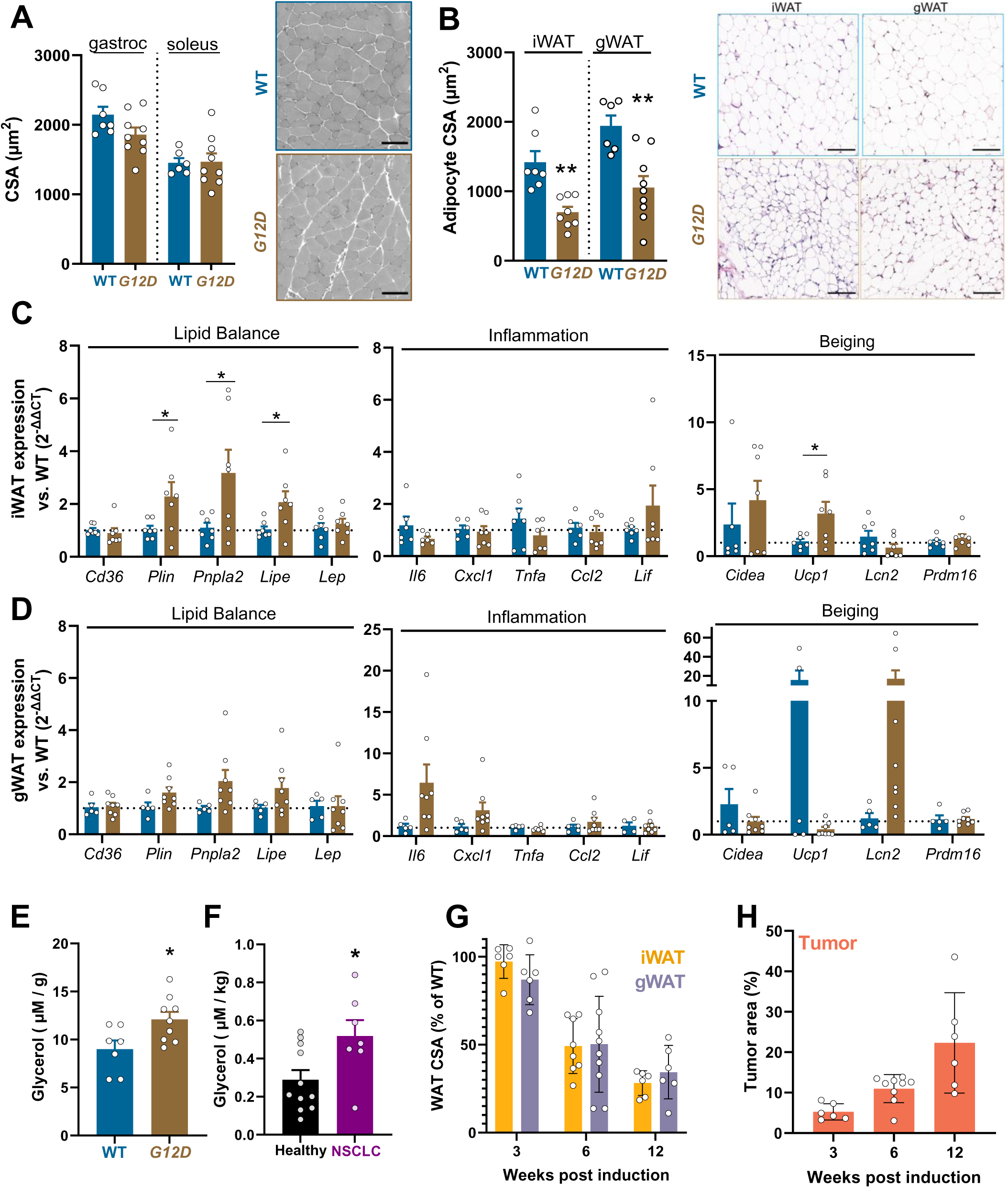
Effect of lung club epithelial specific, inducible Kras^G12D/+^ (G12D) on skeletal muscles and adipose tissues 6 weeks post-induction. Muscle fiber cross-sectional area (CSA) in gastrocnemius and soleus muscles of wild-type (WT; n=7; blue bars) and *G12D* (n=8; gold bars) mice were assessed 6 weeks post tamoxifen injection, with representative images (scale bars=100 µm) (A). Cross-sectional area (CSA) of adipocytes from inguinal (iWAT) and perigonadal (gWAT) white adipose tissue depots in 6-week WT and *G12D* mice, with representative images (scale bars=100 µm) (B). Gene expression in iWAT (C) and gWAT (D) of *G12D* and WT mice for lipolysis, inflammation, and beiging. Glycerol levels in WT and *G12D* mice (E), and in plasma of patients with late-stage, non-small cell lung cancer (NSCLC; n=7; purple bar) compared with healthy controls (Healthy; n=11; black bar) (F). White adipose tissue CSA in *G12D* mice expressed as a percentage of WT CSA at each timepoint (G). Tumor area expressed as a percentage of the total lung area over the 12-week study period (H). Data (A-D) were analyzed Student’s t-test with asterisks indicating * *p*<0.05 and ** *p*<0.01. See also: supplemental figures S7-13.

In keeping with adipocyte wasting, we observed greater circulating glycerol levels in *G12D* mice compared with WT mice (**Fig. 4E**), a marker of increased adipose tissue lipolysis, the process driving adipose wasting in human CC^34^. Patients with NSCLC similarly showed elevated circulating glycerol levels (**Fig. 4F**), suggesting that increased lipolysis in *G12D* mice translate to human lung cancer. Finally, to further discriminate the timing of adipose wasting, we evaluated *G12D* and WT mice 3 weeks post-induction. At 3 weeks, we found no differences in fat pad or organ weight (**Fig S10A**), and no differences in adipocyte or muscle fiber CSA (**Fig S10B-D**). Examining *G12D* adipocyte size over time relative to WT, we found that the progressive reduction in adipocyte size (**Fig 4E**) was inversely associated with increasing tumor burden (**Fig. 4F**). Coupled with the lack of muscle wasting at 3- and 6-weeks, our data suggest that early adipose tissue wasting is important for CC initiation and potentially tumor growth.

Absent differences in energy intake or fecal energy content (Fig S4), and considering that physical activity is not elevated in CC, we explored energetically futile cycles that may underlie elevated energy expenditure. In some models of CC, adipose wasting may be explained by beiging of white adipose tissue ^35, 36^. In *G12D* mice, however, there were no differences in thermogenic gene expression or markers of beiging at 12 weeks post-induction, aside from an upregulation of *Cidea* in gWAT (Fig. 2E-F). In fact, *Ucp1* mRNA expression was lower in iWAT. At 6-weeks post-induction when weight loss begins, there was a modest elevation in mRNA levels of *Ucp1* in iWAT and a trend towards a reduction in eWAT in *G12D* mice (Fig. 4F-G) but no difference in other markers of beiging. To further examine *Ucp1* expression, we measured UCP-1 tissue protein content via immunohistochemical staining ^36^ (**Fig. S11**) and found no differences in UCP-1-positive area (**Fig. S12**), in agreement with work in other mouse models of CC^37^. Finally, whereas UCP-1 expression in other *Kras*-driven lung CC models was dependent on increased circulating IL-6 ^36^, *G12D* mice did not show alterations in circulating IL-6 concentrations (Fig 1E, 3E) or adipose tissue *Il6* mRNA levels (Fig. 2E-F, 4F-G). Thus, there is no evidence for upregulation UCP-1-associated WAT beiging.

To explore other UCP-1-independent energy consuming processes that may contribute to adipose wasting, we evaluated bulk RNA-seq data on adipose for alterations in other energy wasting futile cycles (lipid ^37^ creatine ^38^, Ca^2+^cycling ^39^). Using pathways analysis, with genes involved in these futile cycles included as custom pathways (**Supplemental Table 1**), pathway enrichment analysis showed upregulation of fatty acid metabolism and lipogenesis, including our custom pathway of genes for lipid cycling at 3 weeks (**Fig S9**). The expression of genes for creatine and Ca^2+^ cycling (**Supplemental Table 3**), in contrast, did not differ between groups. Upregulation of pathways of lipid turnover in adipocytes are congruent with increased serum glycerol in *G12D* mice (Fig 4E) and implicate increased lipid cycling as a potential source of the energetic imbalance contributing to CC, in agreement with other models of CC^37^. At 6 weeks post-induction, pathway enrichment analysis showed that pathways for fatty acid metabolism and adipogenesis, including our custom pathway for lipid cycling, were among the top down-regulated genes (**Fig S9**), which may reflect a compensatory adaptation to slow adipose wasting given that considerable atrophy has occurred by 6 weeks (**Figs 4B, S8A, S9**). At this time, neither creatine nor Ca^2+^ cycling pathways were upregulated.

Finally, as tumors may cause liver metabolic reprogramming, we characterized liver phenotypes. Liver weights did not differ at 3 (**Fig S10A**) or 6 weeks (Fig 3D) but were reduced at 12 weeks (Fig 1D). Additionally, we observed a steady increase in liver triglyceride content in *G12D* mice, which became significant by 12 weeks (**Fig S13**), which may result from upregulated adipose tissue lipolysis (Fig 4E and ^40^). Thus, in addition to adipose and skeletal muscle tissue, our model shows evidence for liver metabolic reprogramming ^41–44^, underscoring its utility as a model of multi-organ, systemic metabolic reprogramming in CC ^45^.

### Tissue wasting phenotypes in lung epithelial cell specific Kras^G12D/+^-Stk11 null mice

As patients with lung cancer with mutated *KRAS* lose more weight than patients with wild-type *KRAS*^46^, our model is relevant to human lung tumor oncotypes that predispose to CC. Recently, studies suggest that serine/threonine kinase 11 (*STK11*) mutations are common among patients with NSCLC who report weight loss at diagnosis^47^. Additionally, mice with combined *Kras^G12D/+^-Stk11^-/-^* demonstrate more rapid weight loss^48^ compared with *G12D* mice (Fig. 1C). We therefore hypothesized that lung epithelial-specific *Kras^G12D/+^-Stk11^-/-^*should present a more aggressive form of CC than *Kras^G12D^* alone. To examine this hypothesis, we generated mice with tamoxifen-inducible, club cell specific *Kras^G12D/+^*-*Stk11^-/-^* (**Fig. 5A**). Following tamoxifen induction, *KS* mice exhibited body weight loss over a period of 3.5 weeks (**Fig. 5B**), which diverged from *G12D* and WT 1.5 weeks post-induction, independent of differences in food intake (**Fig. 5C**). More rapid weight loss in *KS* mice was coincident with greater tumor burden compared with *G12D* mice (**Fig. 5D**), and weight loss was related primarily to loss of adipose tissue, as no differences in skeletal muscle or organ weights were observed (**Fig 5E**). *KS* mice had greater circulating levels of IL-6 compared with WT and *G12D* mice and greater CXCL1 compared with WT mice, but no differences in other inflammatory cytokines (**Fig. S14)**. Finally, gastrocnemius muscle fiber CSA was not statistically different among *KS, G12D,* and WT animals (**Fig. 5F**), whereas adipocyte CSA in iWAT was smaller in *KS* compared with *G12D* mice and in gWAT was lower in *KS* and *G12D* compared with WT (**Fig. 5G**). Collectively, these data confirm reports that combined *Kras^G12D/+^-Stk11^-/-^*promotes rapidly progressing CC, show that increasing the number of driver mutations hastens CC development, and indicate that early adipose wasting is maintained across *G12D*-driven models of lung CC with different rates of weight loss.

**Figure 5.**
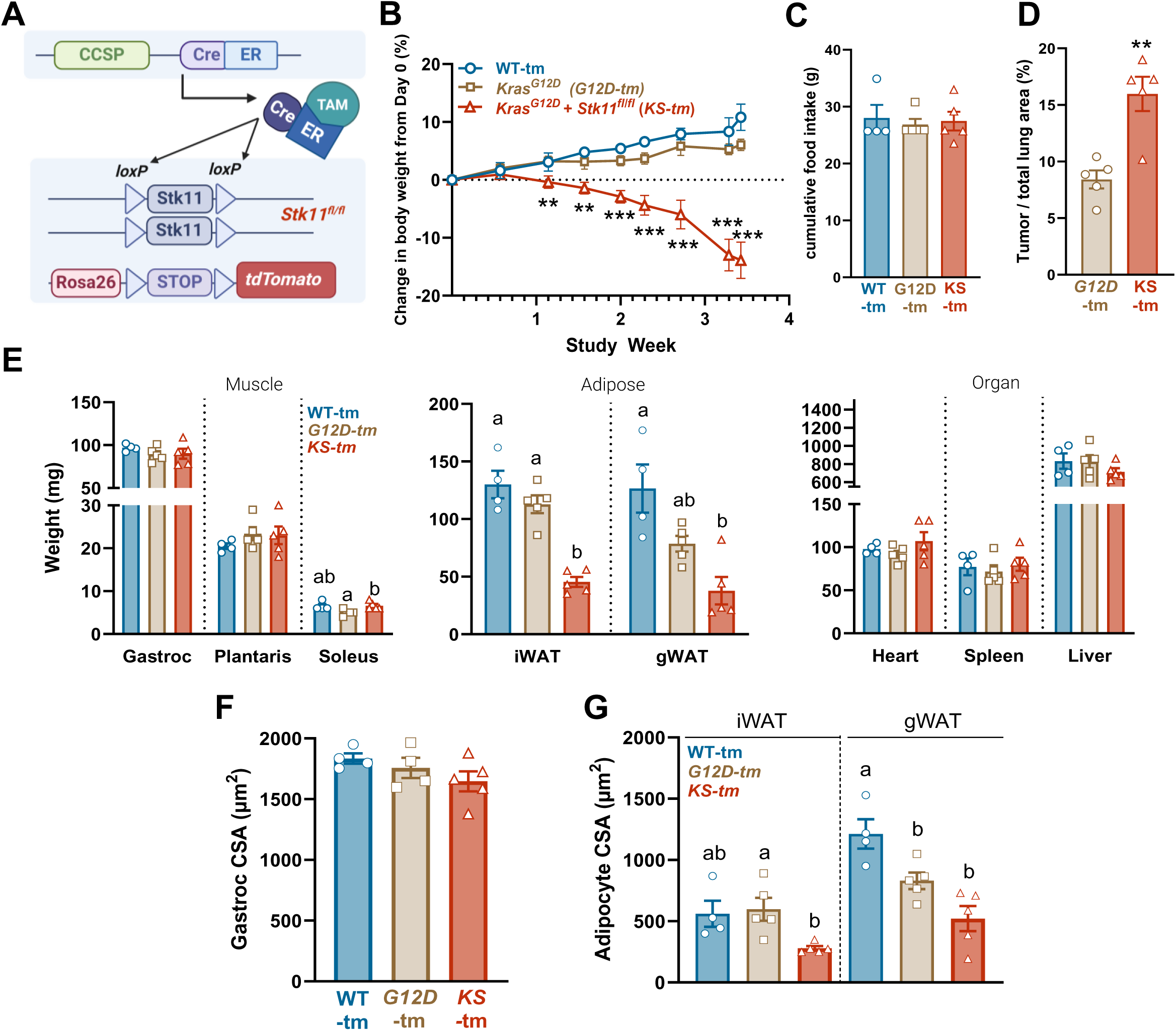
Comparison of lung epithelial cell specific, inducible Kras^G12D/+^ (G12D) and Kras^G12D^/Stk11^-/-^ (KS) mice. *KS-tm* mice harboring tamoxifen (TAM)-inducible lung epithelial cell specific *Kras^G12D^*, deletion of *Stk11/Lkb1* and a *Rosa26-tdTomato (tm)* reporter allele (A). Percent change in body weight (B), cumulative food intake (C) and tumor burden (D) of wild-type (WT-tm; n=4; blue lines/bars/circles), *G12D-tm* (n=5; gold lines/bars/squares) and *KS-tm* (n=5; red lines/bars/triangles) mice in. Muscle, fat pad and organ weights at necropsy (E). Mean gastrocnemius (Gastroc) muscle fiber (F) and iWAT and gWAT adipocyte (G) CSA. Data were analyzed using a repeated measures ANOVA (B), Student’s t-test (D), or a one-way ANOVA (C, E-G) with post hoc comparisons *p*<0.05 group or group X time interaction effects. ** *p* <0.01 different letters by post hoc comparisons. Image created with BioRender.com (A). See also: supplemental figure S14.

### Tumor-derived factors promote CC and adipose tissue wasting

CC is thought to be triggered by factors released from tumors^49^. To examine whether tumor-derived factors initiate adipose tissue wasting, we developed tumor and lung tissue organoids from WT and *G12D* mice, respectively (**Fig. 6A**), from lungs collected at 6-weeks post-induction. Using bulk RNA-seq, we confirmed that the *Kras^G12D^* mutation present *in vivo* was retained in the cultured *G12D* organoids (**Fig. 6B**). Moreover, *G12D* lung tumor organoids showed 1,076 differentially expressed genes (DEG; 541 upregulated and 535 downregulated) compared with WT lung tissue organoids (**Fig. 6C**), with upregulation of numerous oncogenic pathways (**Fig. 6D**). As studies suggest that tumor-derived cytokines contribute to CC ^49–52^, we examined differences between in the expression of inflammatory genes and found 35 DEG, with 23 upregulated and 12 downregulated (**Fig. S15**). None of these DEGs have been implicated in CC, except *Gdf15* (growth differentiation factor-15) ^53^, the expression of which was elevated in *G12D* tumor organoids, with no elevation of GDF-15 in serum from *G12D* mice (Fig. 3E and 1E). Thus, we found no evidence that cytokines from *G12D* tumor organoids may incite CC by increasing systemic inflammation.

**Figure 6.**
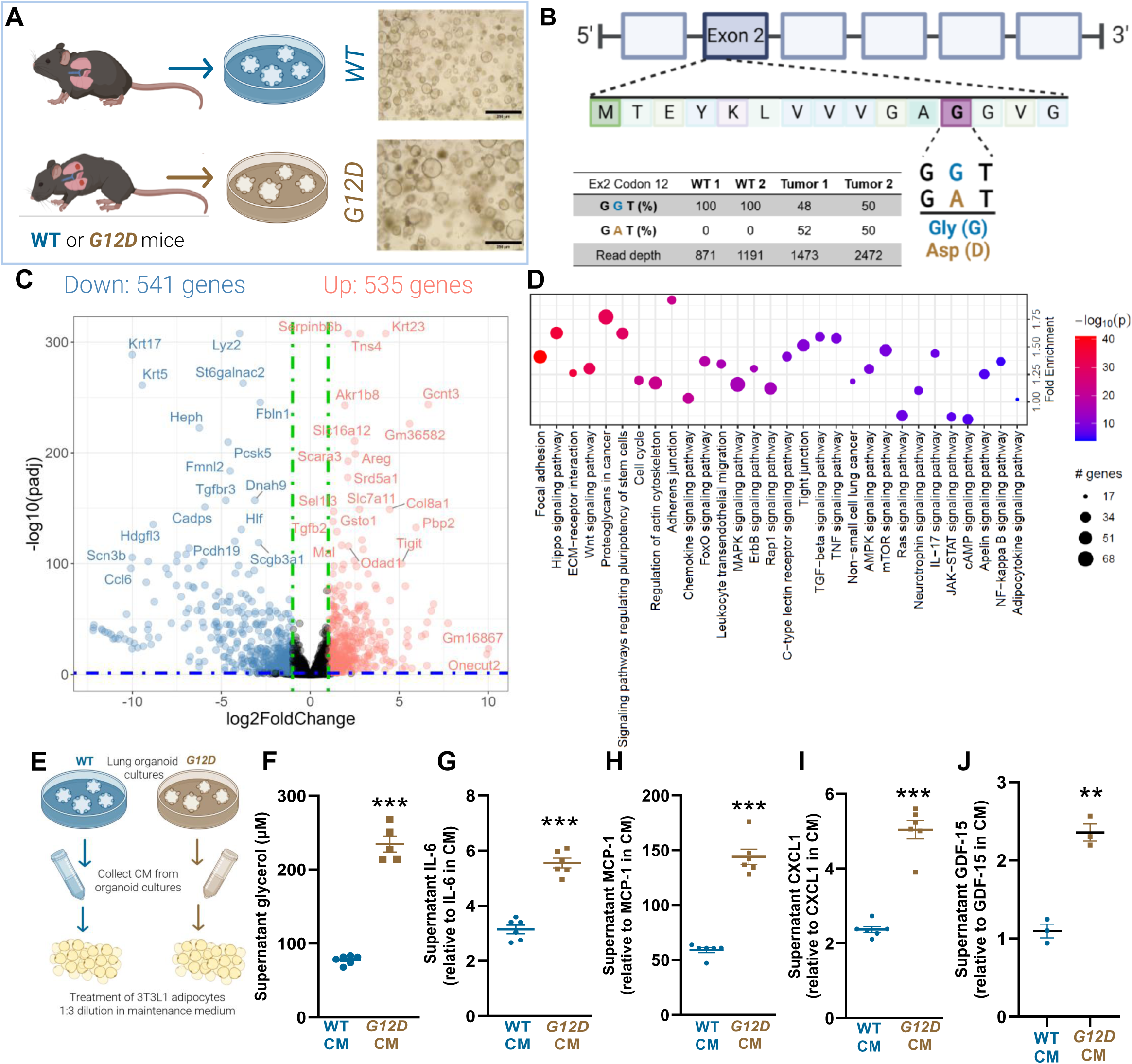
Organoids developed from lung tissue from wild-type (WT) and lung tumors from Kras^G12D/+^ (G12D) mice and their characterization. Lung organoid development with representative images of wild-type (WT; blue) and *G12D* organoids (gold; A; scale bars = 250 μm). RNA-seq was used to validate the *Kras^G12D^* mutation (gold) in lung tumor organoids (B). Volcano plot depicting differentially expressed genes, with red representing upregulated (n=535) and blue representing downregulated (n=541) genes, respectively, in *G12D* compared with WT organoids, with a *p*-value exclusion of 0.05 (C). Pathway analysis depicting upregulated signaling pathways in *G12D* lung tumor compared with WT lung tissue organoids (D). CM was applied (1:3 with adipocyte culture media) to fully differentiated 3T3L1 adipocytes (A). Glycerol levels of supernatant of 3T3L1 adipocytes treated with WT (blue circles) or *G12D* (gold squares) organoid CM for 24 h (F). Supernatant levels of IL-6 (G), MCP-1 (H), CXCL1 (I) and GDF15 (J) from 3T3L1 adipocytes treated for 24 hours with WT or *G12D* organoid CM. 3T3L1 experiments are representative of 2-3 independent replicates. Student’s t-tests were used, with *** *p*<.001. Images created with BioRender.com (A, E). See also: supplemental figures S15-16.

To determine whether tumor-released factors may promote CC in *G12D* mice, we examined whether factors secreted from *G12D* lung tumor organoids alter adipocyte metabolism to promote atrophy by treating murine 3T3L1 adipocytes with conditioned media (CM) from WT tissue or *G12D* tumor organoids (**Fig. 6E**) and assessed lipolysis by measuring media glycerol levels. Treatment of adipocytes with *G12D* tumor organoid CM for 24 hours increased media glycerol levels compared with WT organoid CM (**Fig. 6F**) independent of glycerol levels in *G12D* CM (**Fig. S16A**).

To define how tumor-derived factors increase glycerol release in adipocytes, we evaluated potential mediators. *G12D* CM increased media inflammatory cytokine levels compared with WT CM (**Fig. 6G-I**), independent of cytokine levels in CM (**Fig. S16B-E**). As one of these cytokines, IL-6, has been implicated as a mediator of CC and stimulates lipolysis^54^, we applied *G12D* lung tumor organoid CM that was treated with a neutralizing antibody directed against IL-6 or an isotype control to 3T3-L1 adipocytes. The IL-6 neutralizing antibody reduced IL-6 in the culture media to undetectable levels (**Fig. S16F**), but glycerol levels remained elevated (**Fig. S16G**). Moreover, adding IL-6 to WT lung tissue organoid CM did not increase glycerol release (**Fig. S16F-G**), suggesting that IL-6 is neither necessary nor sufficient to explain increased adipocyte glycerol release.

GDF-15 is slightly increased in *G12D* CM, was recently shown to be increased in circulation of patients experiencing weight loss with recurrence of lung cancer^55^ and may increase adipocyte lipolysis ^56^. To explore whether elevated GDF-15 mediates the increased glycerol release, we treated 3T3-L1 adipocytes with GDF-15 at levels found in *G12D* (2,500 pg/mL) and WT (1,500 pg/mL) organoid CM (**Fig. S16E**). While GDF-15 at both levels increased glycerol above vehicle control, there was no difference in glycerol between the two concentrations, suggesting that GDF-15 likely does not increase adipocyte lipolysis.

Studies have also implicated TLR4 activation in adipose wasting in the LLC model of lung CC^57^. Thus, we evaluated whether TLR4 activation via lipopolysaccharide (LPS) increased glycerol release by 3T3-L1 adipocytes. Activation of TLR4 by LPS was confirmed by robust cytokine release into the media (**Fig. S16I-K**). However, LPS did not increase glycerol release in 3T3-L1 adipocytes (**Fig S16L**), arguing against TLR4 activation as the mediator of *G12D* CM effects on 3T3-L1 adipocytes.

As we observed an increase in adipose inflammation coincident with tissue wasting (Fig. S9) and that macrophages in tissues may drive tissue wasting in CC ^58, 59^, we examined the effects of co-culturing bone marrow-derived macrophages (BMDM) with adipocytes on the cytokine/chemokine response to CM. We found that BMDM enhanced the ability of *G12D* CM to increase MCP-1 and CXCL1 release (**Fig. S16N-O**), but not IL-6 (**Fig S16M**), suggesting that macrophages may recruit circulating immune cells in response to tumor-derived factors to drive local inflammation (Fig S9B and ^30^). Thus, *G12D* tumor organoids release factors that increase adipose tissue lipolysis and may contribute to the development of inflammation in adipose tissue *in vivo* (Fig S9D), phenotypes that parallel adipocyte wasting and inflammation in patients with CC^34, 59, 60^.

Finally, we evaluated whether *G12D* CM could promote wasting of skeletal muscle cells in culture. Treatment of mature, murine C2C12 myotubes ^61^ with *G12D* CM for 72 hours did not reduce myotube diameter compared to WT CM (**Fig. S16P**), in keeping with the lack of effects of CM from 2D cultures of *Kras^G12D^*-transformed murine lung epithelial cells and human lung cancer cell lines^61^.

### Early adipose tissue wasting in G12D mice translates to human patients with lung cancer

To examine whether early changes in adipose tissue in *G12D* mice translate to patients with lung cancer, we carried out a secondary analysis using body composition data collected during the first 3 years of the National Lung Screening Trial (NLST), a longitudinal study examining the efficacy of yearly, low-dose chest computed tomography (CT) scans for early lung cancer diagnosis^62^ (baseline patient characteristics in **Supplemental Table 2**). Using multivariable-adjusted models, females diagnosed with stage 3-4 lung cancer exhibited a reduction in visceral adipose tissue (VAT) cross-sectional area indexed to height (**Fig. 7A**) and an increase in VAT density (**Fig. 7B**; reduced adipocyte size ^63^) compared with controls. In males with stage 3-4 lung cancer SAT density (**Fig 7C**) and skeletal muscle were decreased compared with controls (**Fig. 7D**). Data for non-significant results are shown in **Fig. S17-18**. Our results demonstrate that there is adipose loss prior to diagnosis in patients with lung cancer and a sex dimorphism in these adaptations. Nonetheless, these data show that early adipose tissue adaptations observed in *G12D* mice translates to human lung cancer.

**Figure 7.**
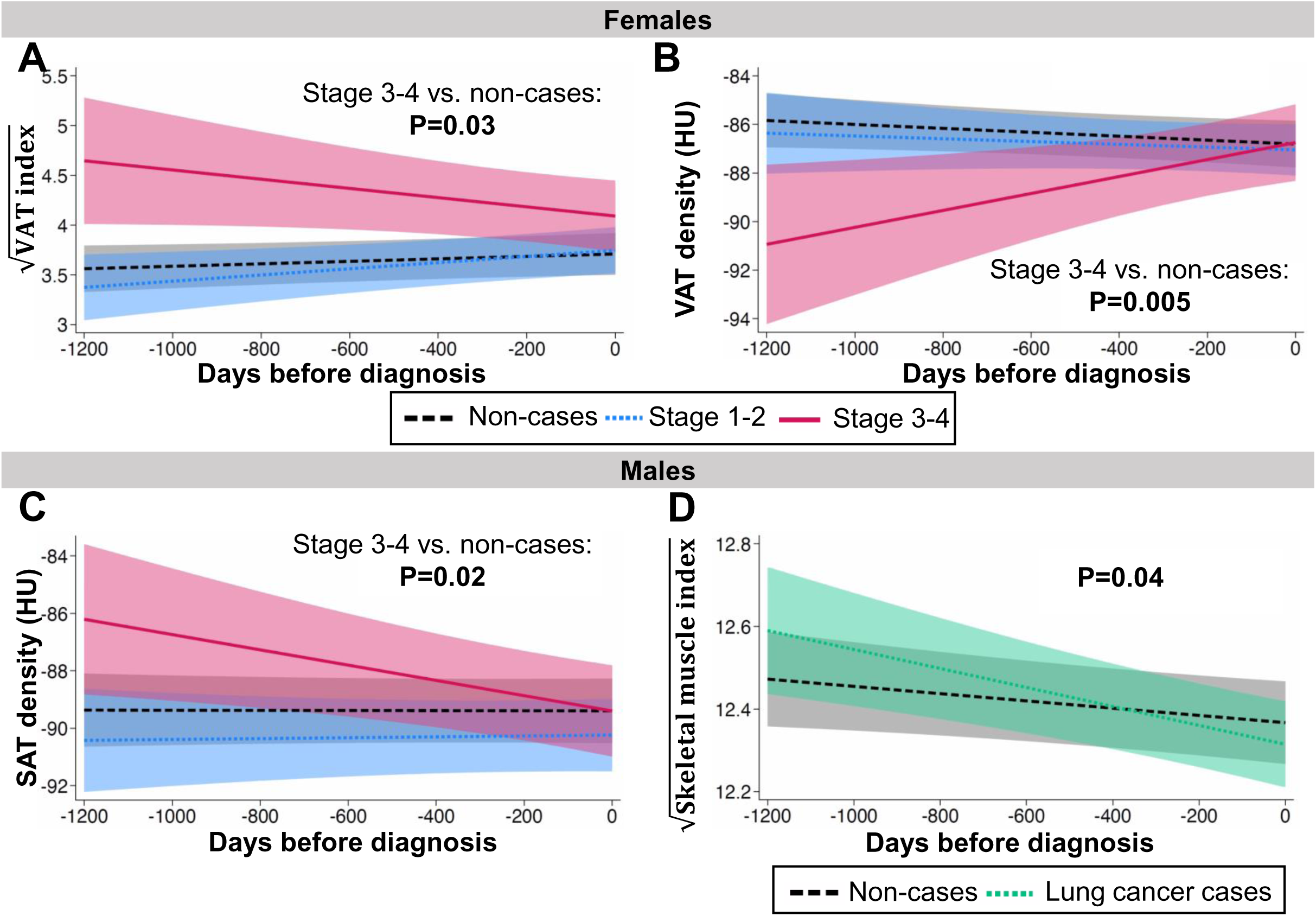
Changes in body composition prior to lung cancer diagnoses among a matched cohort sample of patients in the National Lung Screening Trial (NLST). We conducted secondary analysis of body composition from computed tomography scans collected during the first three years of the NLST using multivariable hierarchical linear models. Modeled data were visualized by plotting marginal predicted values across the range of time values by exposure group and stratified by sex, and we present 95% confidence intervals as shaded regions. Data are shown for non-cases (black lines) and patients diagnosed with stage 1-2 (blue) or stage 3-4 lung cancer (A) when stratified by diagnosis and stage or by non-cases (black lines) and lung cancer cases (green lines) when stratified by diagnosis only. Visceral adipose tissue (VAT) area indexed to height transformed via square root to meet normality (A) and density in Hounsfield units (HU), measured by tissue attenuation (B). Subcutaneous adipose tissue (SAT) density, measured by tissue attenuation (C). Skeletal muscle area indexed to height in non-cases and lung cancer cases and transformed via square root to meet normality (D). *P*-values and comparison groups are listed in each respective figure. See also: supplemental figures S17-18.

## Discussion

We report the development of a new model of lung CC using mice with a lung epithelial cell-specific, tamoxifen-inducible *Kras^G12D^* mutation. *G12D* mice exhibit more physiologically relevant tumors with respect to their anatomic location and burden, a slower time course of CC initiation and progression, and peripheral tissue and tumor morphological, mutational, transcriptomic, and metabolic characteristics that parallel CC in patients with lung cancer.

Moreover, adipose tissue wasting drove CC initiation, could be incited by *G12D* lung tumor released factors, was inversely related to tumor burden, was apparent across different *Kras*- driven lung CC models with differing mutational burdens and induction approaches, and translated to human patients, highlighting an early role for tumor-induced metabolic reprogramming of adipose tissue wasting in CC.

### Tumor location, burden, and mutations

Most studies of CC have employed heterotopic murine models, in which tumor cells are injected into the flank, producing a tumor akin to a metastatic lesion but in a microenvironment (*i.e.*, subcutaneous) that is atypical outside of melanoma. Recent studies have advanced this approach by administering cancer cells with combined *Kras* and *Tp53* mutations into the vasculature or lungs to seed tumors in the lung ^64, 65^. In *G12D* mice, the transgene is targeted to lung epithelial cells, and tumors originate in the terminal bronchioles, the anatomic location of most lung tumors in humans ^66^. Moreover, at the initiation of weight loss (6 weeks post-induction), tumor burden in *G12D* mice is small (<<1% of body weight) relative to other models, better reflecting tumor burden in patients. An additional advantage of *G12D* mice is that *KRAS* is the most frequently mutated oncogene ortholog in human NSCLC ^67^, the majority of patients with lung cancer have elevated *RAS* pathway activation ^68^ and those with mutated *KRAS* experience greater weight loss ^46^. Building on recent reports suggesting that *STK11* loss-of-function amplifies lung CC in humans ^47^, our data demonstrate that mice with combined *Kras^G12D^/Stk11^-/-^* develop a more rapidly progressive CC than *Kras^G12D^* alone, and comparable to the timeframe of recently developed models using cells with *Kras/Tp53* mutations ^64, 65^. Thus, our approach of using inducible, epithelial cell specific expression of transgenes provides a useful platform to study human lung tumor oncotypes that predispose to CC and offers a model that approximates tumor characteristics in human lung CC with respect to anatomic location, burden, and mutations that predispose to CC. Based on our results showing potentially different tissue wasting phenotypes when *Kras^G12D^* is expressed in different lung cell types (club vs. AT2 cells), further studies are needed to evaluate cell- and driver mutation-specific variation in CC and how this relates to CC in patients with different tumor oncotypes ^46, 47^.

### CC initiation and progression

While recent lung CC models have made advances in localizing tumors to the lungs ^64, 65^, they show rapid onset and progression of weight loss, which does not reflect what occurs in patients. In contrast, the initiation and progression of weight loss in *G12D* mice is slower, with an ∼4-6-week latency until weight loss begins, followed by progressive weight loss until 12 weeks post-induction. The relatively slow rate of progression of weight loss in *G12D* mice (∼1.9%/week) more closely approximates that in patients with lung cancer ^69^ and offers a comparatively longer time to intervene on CC with therapeutics (∼6-8 weeks).

### Early adipose tissue wasting

Exploiting the protracted time course of the *G12D* model to examine the temporal context of body composition changes in CC, we report early wasting of adipose tissue in *G12D* mice. Considering the relative contribution of adipose tissue to body weight in adult C57BL/6J mice^70^ and our observed reductions at 6 weeks post-induction (>50%), loss of adipose could explain the majority of body weight loss. Reinforcing these data, we found early adipose tissue wasting in mice with combined *Kras^G12D/+^-Stk11^-/-^*, which exhibit more rapidly progressive weight loss than *G12D* mice, and in *Kras*-driven lung cancer induced via adenoviral Cre recombinase, suggesting that early adipose tissue wasting remains a defining characteristic of *Kras-*driven lung CC across models that differ in mutational burden, time course, and induction techniques.

Factors released from *G12D* tumors may drive early adipose tissue wasting, as we found that cultured adipocytes treated with *G12D* lung tumor organoid CM release greater amounts of glycerol into the media, a proxy of lipolysis, compared with adipocytes treated with WT lung tissue organoid CM. Similar effects were evident *in vivo*, as circulating glycerol levels were increased in *G12D* mice at 6 weeks post-induction, when weight loss begins. In fact, trends towards elevated circulating glycerol remained at 12 weeks post-induction, along with increased circulating markers of fat availability/metabolism. Moreover, we found elevated glycerol levels in patients with NSCLC compared with non-cancer controls. As increased lipolysis contributes to adipose wasting in other animal models of CC^37^ and humans ^34^, these data suggest that tumor-released factors may drive elevated adipose tissue lipolysis in *G12D* mice.

The importance of early adipose tissue wasting is apparent in recent prospective data in patients with pancreatic cancer, which showed that the relative loss of adipose tissue was twice that of skeletal muscle prior to diagnosis and was a stronger predictor of body weight loss ^71^. Moreover, longitudinal studies show that adipose wasting constitutes the majority of weight loss in patients with CC ^72^. Patients with lung cancer lose body weight prior to diagnosis ^69^ but, based on our recent systematic review, there are no longitudinal data examining early body composition changes^73^. To address this lack of data, using AI-based approaches ^74, 75^, we conducted a secondary analysis of data from the NLST ^76^. Both male and female patients with advanced-stage disease experience changes in adipose tissue size and composition prior to lung cancer diagnosis, with females experiencing a decrease in VAT size and an increase in density. Sex differences in the anatomic localization of adipose in the thoracic compartment, where CT scans were conducted, make it challenging to draw conclusions about biological sex-specific changes^77^ and further studies evaluating other adipose depots and/or whole-body adiposity are needed. Nonetheless, these data suggest that early adipose tissue wasting in *G12D* mice translates to human patients.

Early adipose tissue wasting in CC may have pathological consequences, as lipid uptake is required to support cancer cell proliferation under certain conditions^78^ and elevated or reduced availability of lipid substrates in conditions of high-fat feeding or caloric restriction, respectively, increases or decreases proliferation^79^. Further suggesting a role for adipose tissue substrates in cancer progression, we found an inverse association between adipocyte wasting and tumor burden in *G12D* mice. The modulatory role of endogenous lipid substrates on tumor growth is an important topic given the current epidemic of obesity, and our results underscore the utility of the *G12D* model to evaluate a potentially novel role for CC-induced adipose tissue wasting in cancer progression.

### Limitations of the Study

Several limitations to our study deserve discussion. First, tumors in *G12D* mice were constrained to the lung, limiting the ability of this model to study the impact of tumor metastases on CC. Second, organoids do not reproduce all cell types and the physiology of tumors in the lung, nor does use of their CM in cultured adipocytes reproduce the complex array of cells, organs and tissues *in vivo*. Similarly, because organoids are developed from whole lung homogenates, we cannot evaluate how lesion heterogeneity contributes to CC. Third, as the NLST does not contain information on tumor oncotype, it is unclear if body composition changes reflect the effect of mutant *KRAS*. However, 84% of lung adenocarcinomas have elevated *RAS* pathway activation^68^, suggesting that tumors in most NLST patients likely exhibit increased *RAS* pathway activation. Finally, we did not identify the molecule or molecules that are necessary and sufficient for CC in *G12D* mice. Considering the protracted nature of CC and differential time course of adipose and skeletal muscle wasting, these could vary by tissue type, along the continuum of CC, and/or involve complex interactions with other organs and tissues. Further work will need to identify the complex tumor-host interactions that drive lung CC in this and other clinically representative models to derive therapies that can alleviate CC.

### Conclusions

Our data show that the *G12D* mouse lung cancer model phenocopies tissue, cellular, mutational, transcriptomic, and metabolic characteristics of human CC. Additionally, because of the protracted time course for initiation and progression of CC relative to other pre-clinical lung CC models, *G12D* mice are well-suited to identify mechanisms initiating CC, as well as testing preventative and remediative therapies. Using this model, we identify adipose tissue wasting as an early event in *G12D* and *Kras^G12D^-Stk11^-/-^* mouse models of lung CC that may be mediated by tumor-released factors. The translational relevance of this model is supported by our data from a large, prospective trial, where we observed that adipose wasting prior to lung cancer diagnosis. Together, these findings underscore the utility of this model to study the mechanisms of CC and to the role of early adipose tissue wasting in CC and cancer progression.

## Acknowledgements

This study was funded by NIH R01 AR065826 (MJT), R21 CA283492 (MJT), T32 HL076122 (MEP), R21 CA280266 (DJS), R01 CA273238 (YJ-H), R01 CA253923 (BAL), and the University of Vermont Cancer Center. Imaging work was performed at the Microscopy Imaging Center at the University of Vermont (RRID# SCR_018821). The Graphical Abstract was created in BioRender. Snoke, D. (2025) https://BioRender.com/zxbk8g3. A UVM College of Medicine Shared Instrumentation Award to D.T. supported the purchase of the Leica-Aperio VERSA whole slide imaging system used for image acquisition. The University of Colorado SOM Metabolomics Core is supported in part by the University of Colorado Cancer Center via NIH P30 CA046934 and the University of Michigan Animal Phenotyping Core via grants U2CDK110768, KD020572, and DK089503. Funding in support of the RNA-seq was provided by a Zymo Impact Initiative Grant (DBS). The authors thank Vikas Anathy and Zoe Mark for the machinery and technical support needed to measure oxygen saturation.

## Author Contributions

Conceptualization: MJT, DBS; Methodology: DBS, MEP, DM, DJS, RA, YJH, KX; Validation: DBS ; Formal analysis: DBS, DM, JSD, TPA; Investigation: DBS, JLA, JV, AEB, RA, SML, HS, ERB, SCJH, MMM, DS, JAR; Resources: MJT, MEP, JV, DM, DJS, YJH, AD, BAL, KLS; Data Curation: DBS, HS, ERB, SCJH, SML, DS, KX; Writing-original draft preparation: DBS, MJT; Writing-review and editing: all authors; Visualization: DBS, TPA, JSD; Supervision: MJT, MEP, DJS, DBS; Project administration: DBS and MJT; Funding acquisition: MJT, MEP, YJ-H and DBS.

## Methods

### Resources table

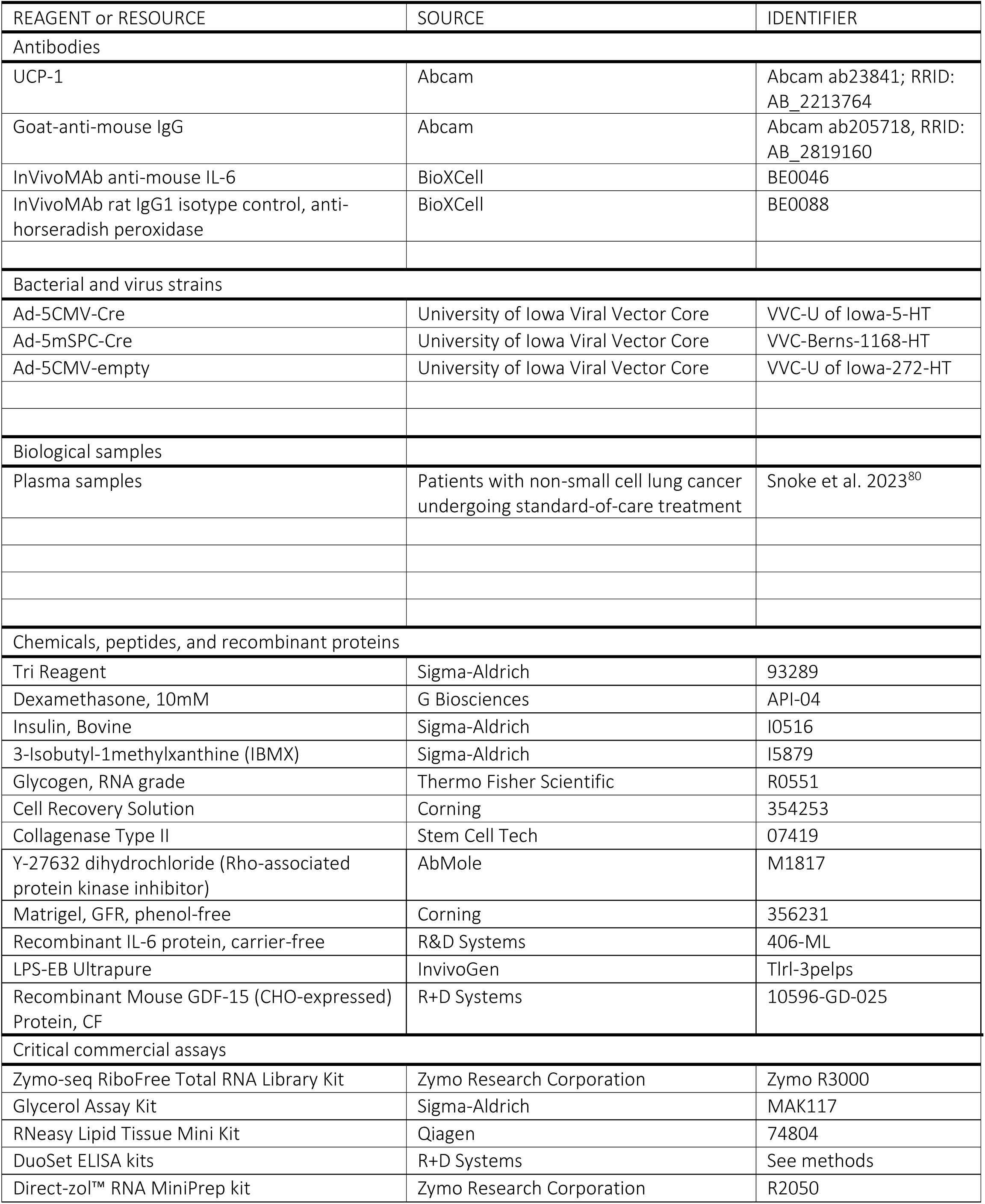
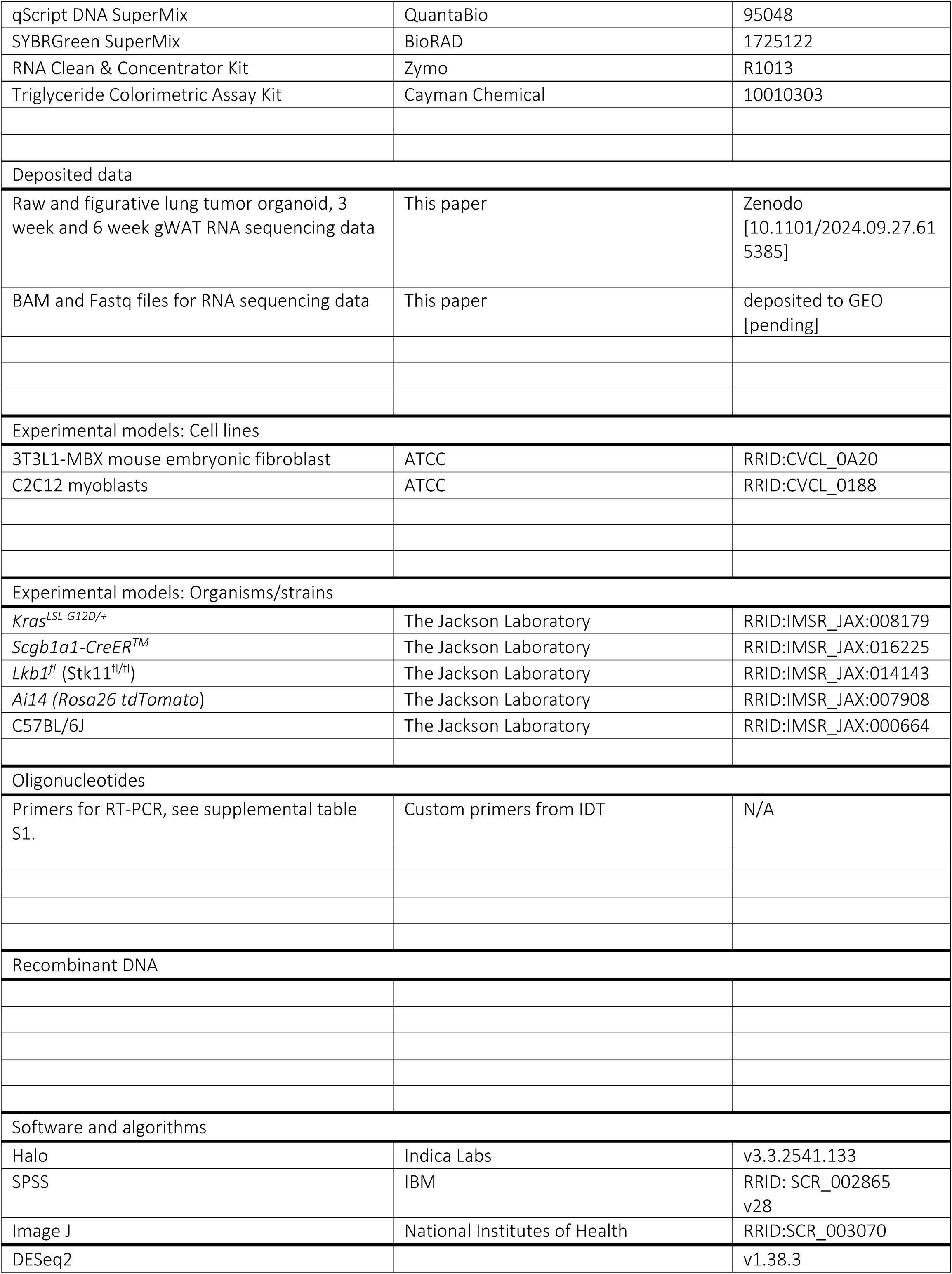
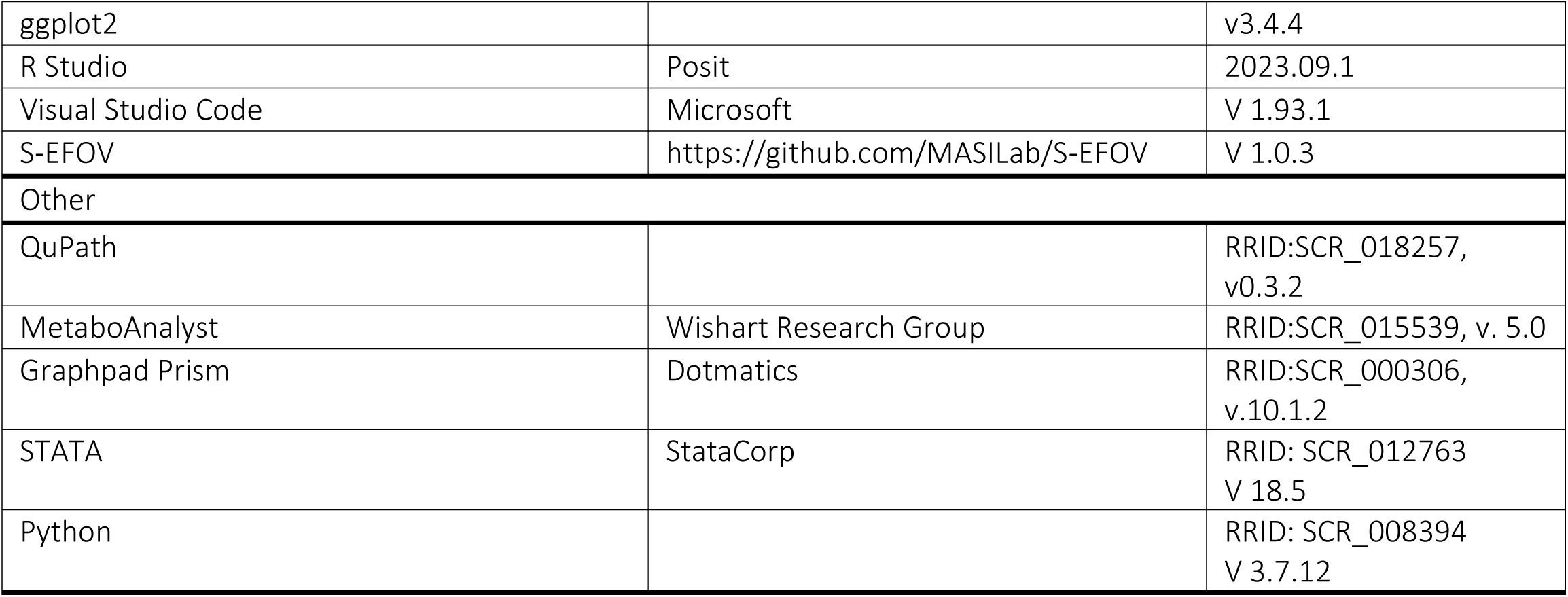

| REAGENT or RESOURCE | SOURCE | IDENTIFIER |
| --- | --- | --- |
| **Antibodies** |  |  |
| UCP-1 | Abcam | Abcam ab23841; RRID: AB_2213764 |
| Goat-anti-mouse IgG | Abcam | Abcam ab205718, RRID: AB_2819160 |
| InVivoMAb anti-mouse IL-6 | BioXCell | BE0046 |
| InVivoMAb rat IgG1 isotype control, anti-horseradish peroxidase | BioXCell | BE0088 |
| **Bacterial and virus strains** |  |  |
| Ad-5CMV-Cre | University of Iowa Viral Vector Core | VVC-U of Iowa-5-HT |
| Ad-5mSPC-Cre | University of Iowa Viral Vector Core | VVC-Berns-1168-HT |
| Ad-5CMV-empty | University of Iowa Viral Vector Core | VVC-U of Iowa-272-HT |
| **Biological samples** |  |  |
| Plasma samples | Patients with non-small cell lung cancer undergoing standard-of-care treatment | Snoke et al. 2023<sup>80</sup> |
| **Chemicals, peptides, and recombinant proteins** |  |  |
| Tri Reagent | Sigma-Aldrich | 93289 |
| Dexamethasone, 10mM | G Biosciences | API-04 |
| Insulin, Bovine | Sigma-Aldrich | I0516 |
| 3-Isobutyl-1methylxanthine (IBMX) | Sigma-Aldrich | I5879 |
| Glycogen, RNA grade | Thermo Fisher Scientific | R0551 |
| Cell Recovery Solution | Corning | 354253 |
| Collagenase Type II | Stem Cell Tech | 07419 |
| Y-27632 dihydrochloride (Rho-associated protein kinase inhibitor) | AbMole | M1817 |
| Matrigel, GFR, phenol-free | Corning | 356231 |
| Recombinant IL-6 protein, carrier-free | R&D Systems | 406-ML |
| LPS-EB Ultrapure | InvivoGen | Tlrl-3pelps |
| Recombinant Mouse GDF-15 (CHO-expressed) Protein, CF | R+D Systems | 10596-GD-025 |
| **Critical commercial assays** |  |  |
| Zymo-seq RiboFree Total RNA Library Kit | Zymo Research Corporation | Zymo R3000 |
| Glycerol Assay Kit | Sigma-Aldrich | MAK117 |
| RNeasy Lipid Tissue Mini Kit | Qiagen | 74804 |
| DuoSet ELISA kits | R+D Systems | See methods |
| Direct-zol™ RNA MiniPrep kit | Zymo Research Corporation | R2050 |
| qScript DNA SuperMix | QuantaBio | 95048 |
| SYBRGreen SuperMix | BioRAD | 1725122 |
| RNA Clean & Concentrator Kit | Zymo | R1013 |
| Triglyceride Colorimetric Assay Kit | Cayman Chemical | 10010303 |
| Deposited data |  |  |
| Raw and figurative lung tumor organoid, 3 week and 6 week gWAT RNA sequencing data | This paper | Zenodo<br>[10.1101/2024.09.27.615385] |
| BAM and Fastq files for RNA sequencing data | This paper | deposited to GEO<br>[pending] |
| Experimental models: Cell lines |  |  |
| 3T3L1-MBX mouse embryonic fibroblast | ATCC | RRID:CVCL_0A20 |
| C2C12 myoblasts | ATCC | RRID:CVCL_0188 |
| Experimental models: Organisms/strains |  |  |
| *Kras*<sup>LSL-G12D/+</sup> | The Jackson Laboratory | RRID:IMSR_JAX:008179 |
| *Scgb1a1-CreER*<sup>TM</sup> | The Jackson Laboratory | RRID:IMSR_JAX:016225 |
| *Lkb1*<sup>fl</sup> (Stk11<sup>fl/fl</sup>) | The Jackson Laboratory | RRID:IMSR_JAX:014143 |
| *Ai14 (Rosa26 tdTomato)* | The Jackson Laboratory | RRID:IMSR_JAX:007908 |
| C57BL/6J | The Jackson Laboratory | RRID:IMSR_JAX:000664 |
| Oligonucleotides |  |  |
| Primers for RT-PCR, see supplemental table S1. | Custom primers from IDT | N/A |
| Recombinant DNA |  |  |
| Software and algorithms |  |  |
| Halo | Indica Labs | v3.3.2541.133 |
| SPSS | IBM | RRID: SCR_002865<br>v28 |
| Image J | National Institutes of Health | RRID:SCR_003070 |
| DESeq2 |  | v1.38.3 |
| ggplot2 |  | v3.4.4 |
| R Studio | Posit | 2023.09.1 |
| Visual Studio Code | Microsoft | V 1.93.1 |
| S-EFOV | https://github.com/MASILab/S-EFOV | V 1.0.3 |
| Other |  |  |
| QuPath |  | RRID:SCR_018257, v0.3.2 |
| MetaboAnalyst | Wishart Research Group | RRID:SCR_015539, v. 5.0 |
| Graphpad Prism | Dotmatics | RRID:SCR_000306, v.10.1.2 |
| STATA | StataCorp | RRID: SCR_012763 V 18.5 |
| Python |  | RRID: SCR_008394 V 3.7.12 |

### Resource availability

#### Lead contact

Further information and requests for resources and reagents should be directed to and will be fulfilled by the lead contact, Michael Toth.

### Materials availability

#### Data and code availability

- RNA-seq raw read counts will be available via the Zenodo online repository.
- Original code used to generate figurative data will be available in the Zenodo repository and from the lead contact upon request.
- Any additional information required to reanalyze the data reported in this work is available from the lead contact upon request.

### Experimental model and study participant details

#### Animals

All animal experiments were approved by the University of Vermont Institutional Animal Care and Use Committee, in compliance with the National Institutes of Health Guidelines for Use and Care of Laboratory Animals. Mice were housed with sex-matched litter mates and given *ad libitum* access to standard chow and water, except for a subset of mice that were housed singly to monitor food intake as specified. Animals were allocated by genotype.

#### Club cell specific, inducible Kras^G12D/+^ (G12D) mice

*G12D* mice (*Scgb1a1-CreER^TM^-Kras^LSL-G12D/+^*) were generated by crossing *Scgb1a1-CreER^TM^* mice with *Kras^LSL-G12D/+^* mice and were maintained on a mixed background. At 4-5 months of age, *G12D* or wild-type *Scgb1a1-CreER^TM^*(WT) littermates were administered five consecutive daily intraperitoneal (IP) injections of 20 mg/ml tamoxifen (Sigma-Aldrich, St. Louis, MO, USA) dissolved in corn oil at a dose of 1 mg tamoxifen/10 g body weight. Body weight was monitored 2-3 times weekly and food intake was measured in a sub-group of individually housed mice for 6 weeks. At 6- or 12-weeks post-tamoxifen injections, mice were euthanized by cervical dislocation under pentobarbital anesthesia. Blood was collected by cardiac puncture in serum separator tubes and centrifuged to isolate serum, aliquoted, and stored at −80°C. Skeletal muscles, adipose tissues, heart, liver, and spleen were weighed, flash-frozen in liquid nitrogen, and stored at −80°C. Lungs were collected for histology or organoid development. One gastrocnemius, soleus and tibialis anterior muscle were mounted in 7% gum tragacanth, flash-frozen in liquid nitrogen-cooled isopentane, and stored at −80°C. One inguinal (iWAT) and one perigonadal (gWAT) fat pad were fixed in 10% formalin and paraffin-embedded.

#### Kras^G12D/+^ -Stk11^-/-^ Mice

*G12D* mice were crossed with *Lkb1^fl/fl^* (*Stk11^fl/fl^)*^81^ and Ai14 (*Rosa26-tdTomato*; TD) reporter mice^82^ to generate *Kras^G12D/+^*-TD (*G12D)*, *Kras^G12D/+^*-*Stk11^-/-^*-TD (*KS)* and wild-type *Scgb1a1-CreER^TM^-TD* (WT; *Kras^+/+^*) littermate control mice. At 8-10 weeks of age, *G12D, KS*, and WT mice were injected (IP) for five consecutive days with tamoxifen (Sigma-Aldrich, St. Louis, MO, USA) dissolved in corn oil as detailed for *G12D* mice. Body weight and food intake were monitored thrice weekly for 3.5 weeks, after which mice were euthanized, and blood and tissues were collected similar to *G12D* mice.

#### Adeno-Cre and Adeno-5-CMV-Cre mice

Mice harboring *Kras^LSL-G12D^* (JAX 008179) and *Rosa26^LSL-tdTomato^* (Ai14, JAX 007914) alleles were used. Cre recombinase was introduced via oropharyngeal administration of 5 x 10^7^ pfu/mouse of adenoviral Cre (Ad5-CMV-Cre or Ad5mSPC-Cre; University of Iowa Viral Vector Core) diluted in 1X PBS to activate recombination in lung epithelial cells. Initially, mice were monitored once a week and weighed once every other week. At 12-16 weeks post-infection, mice were euthanized by IP injection of 200 mg/kg of sodium pentobarbital and tissues collected for analyses.

#### Lung Tumor Organoids

Tumor or lung tissue from *G12D* or *WT* mice, respectively, were excised 6 weeks post-tamoxifen injection, and organoid cultures were developed, as described ^83^. Briefly, lung tumors or tissue were excised and minced in high glucose GlutaMAX DMEM containing 0.1% BSA and 1% penicillin-streptomycin (D-BSA). Minced tissue was incubated with 500 μL collagenase type II (20 mg/mL) and 10 μL of Rho-associated protein kinase inhibitor (10 μM) on an orbital shaker at 140 rpm for 1-2 hours at 37°C. Fetal bovine serum (FBS) was added to halt collagenase digestion and tissues were further homogenized by vigorous pipetting. The tissue suspension was strained through a 100 µM filter, centrifuged at 8°C, the supernatant aspirated leaving ∼0.5 mL, and the pellets resuspended. Tissue pellets were washed twice in D-BSA and centrifuged at 8° C, and tissue pellets were resuspended in Matrigel (200 μL for a 50 μL pellet). The tissue suspension (50 μL) was seeded into each well of a 6-well plate in ∼10 μL drops. Maintenance of the oncogenic phenotype was confirmed by assessment of the single nucleotide variant (c.35G>A) in the *Kras* allele via RNA-seq. For subsequent *in vitro* studies, conditioned medium (CM) from *G12D* or WT organoids were collected over a 3-day period after organoid passaging. CM was frozen at −80°C in aliquots for *in vitro* studies. For cell treatment, CM was thawed on ice directly prior to 25% dilution (1:3) in cell culture maintenance medium.

#### Cells

3T3-L1-MBX mouse embryonic fibroblasts were cultured and differentiated, as described^84^. For some experiments, cells were treated with recombinant LPS, IL-6, or GDF-15 for a period of 24 hours and supernatants assessed by ELISA.

C2C12 myoblasts were cultured as previously described ^61^. On Day 7 post-differentiation, myotubes were treated with 25% (1:3) WT or *G12D* organoid CM in maintenance medium, replenished daily, for a period of 72 hours. Myotube diameter was assessed using digital images taken at 4x magnification. An average of 5 diameters were taken for each myotube of n=5-25 myotubes per field from n=4 random fields were measured using ImageJ software by an assessor blinded to treatment status.

All cells were used prior to passage 10 and all experiments were independently replicated 2-3 times.

#### Human volunteers

Patients recently diagnosed with non-small cell lung adenocarcinoma (NSCLC; n=11; 6 male, 5 female) and age- and sex-matched healthy controls (HC; n=11; 6 male, 5 female) were recruited, as described ^80^. After an overnight fast, patients with NSCLC receiving evidence-based anti-cancer therapy (82 ± 4 days) and matched controls underwent a fasting blood draw, and plasma was isolated for measurement of glycerol levels. The study was approved by the University of Vermont Committee on Research in the Medical Sciences and written informed consent was obtained from each volunteer before participation. Other data collected from these study participants are published in our recent clinical work^80^.

To understand early, pre-diagnosis changes in muscle and adipose in patients with lung cancer, we conducted a secondary analysis using body composition data collected during the first 3 years of the NLST (ClinicalTrials.gov identifier NCT00047385) –a large, longitudinal study examining the utility of yearly low-dose, helical computed tomography scans vs. standard chest x-ray for earl lung cancer diagnosis. Participants in the original study were 55-74 years of age with a cigarette smoking history of at least 30 pack-years who either smoked currently or had quit smoking within the past 15 years. The computed tomography (CT) arm of the study included 26,722 participants who were invited for, and 25,139 participants who completed CT screening examinations at 1-year intervals over the first 3 years, with baseline screening examinations performed after randomization^76^. Additional details of inclusion and exclusion criteria have been previously described^76^. The exposed group for our analyses included n=436 lung cancer patients with ≥2 CT scans before diagnosis. For each exposed participant, we calculated the days elapsed between their most recent scan and the date of lung cancer diagnosis. To each exposed participant, we matched one participant who was alive and without a lung cancer diagnosis (*i.e.*, unexposed) on the corresponding number of days since their last scan (n=436).Baseline group characteristics for lung cancer cases and controls are listed in Supplemental Table S2. Body composition was assessed using AI analysis of lung screening computed tomography scans, as described previously^74, 75^. We fit crude and multivariable-adjusted hierarchical linear models to estimate changes in adipose tissue and muscle depot size and density, with adjustment for race/ethnicity, BMI, and age, as described in the Statistics section of the Star Methods. Because of sex differences in body composition in the anatomical region examined (thoracic), all analyses were stratified by biological sex

#### Definitions of Analytic Variables from National Lung Screening Trial Data

Our main exposure definition was lung cancer cases vs. non-cases. We further explored parsing lung cancer cases by stage at diagnosis, yielding an exposure definition of stage 3-4, stage 1-2, and non-cases. Following visual evaluation of body composition variable distributions, we applied either log or square root transformations to skeletal muscle index, SAT index, VAT index, and VAT area to improve normality before use in regression models. Patient age was categorized for descriptive purposes but modeled in continuous form. The time scale for longitudinal measurements was calculated as the difference in date of lung cancer diagnosis (or corresponding date for matched non-cases) and date of each CT screening. Race and ethnicity were left in fine categories for descriptive analysis, but sparsity in most categories required simplification to “White, non-Hispanic or Latino” and “Other” for regression modeling. We calculated body mass index (BMI) from baseline weight and height and placed into standard categories for descriptive analysis; regression models used continuous BMI.

### Method details

#### Fecal energy content

During Week 6 post-tamoxifen injection, food intake and fecal samples were collected over a 72-hour period to calculate total energy intake and fecal energy content, as previously described. Within this 72-hour period, fecal samples were collected every 24 hours and immediately frozen at −80C. Samples were dried, powdered, and mixed with wheat flour of pre-measured energy content (10% fecal sample). Samples were shaped into pellets and in duplicate subjected to complete oxidation under O_2_ in a bomb calorimeter (6200 isoperibol bomb calorimeter, Parr) and calorie content and digestible energy content were calculated.

#### Tumor burden

H&E sections from formalin-fixed, paraffin-embedded lung tissues were imaged and analyzed by a board-certified anatomic pathologist (DJS) blinded to the identity of the specimens. Qupath was used to manually annotate tumor vs normal lung tissue and assess % tumor burden in μm^2^ and expressed relative to total lung area (%). The presence of adenocarcinomas was confirmed by a second board-certified anatomic pathologist who was not a member of the research team.

#### Enzyme-linked immunoassay (ELISA)

For organoid CM and cell supernatant, levels of IL-6 (DY-406), MCP-1 (DY-479), CXCL1/KC (DY-453) were quantified using DuoSet ELISA kits, according to the manufacturer’s protocol. DuoSet ELISA Kits were also used to measure TNFα (DY-410) and IL-6 Rα (DY1830) in serum, according to the manufacturer’s protocol. Absorbance was read at 570 nm and background was removed using 450 nm absorbance on a Synergy HTX plate reader and Gen5 software (BioTek Inc, Winooski, VT, USA). Mouse serum cytokines were analyzed by a custom Luminex-based multiplex assay (R&D Systems) for mouse CXCL1, GDF-15, TNFα, IL-6, CXCL10, IL-1β, IL-7, IL-10, TNFR-II, IL-12, IL-17, IL-18, LIF, CXCL10, and FGF21, according to manufacturer instructions. In some instances, cytokines produced in response to *G12D* organoid CM treatment were expressed as the fold change compared to WT organoid CM treatment to account for variability in cytokine production across experimental replicates.

#### Muscle fiber cross-sectional area (CSA)

Transverse cryosections (10 µm) of gastrocnemius and soleus muscles were stained with hematoxylin and eosin, whole slides were scanned (Leica-Aperio VERSA 8, Leica Biosystems, Deerfield, IL), and 2-3 representative 500 µm x 500 µm images from each of two representative levels along the length of the muscle were chosen to quantify the mean muscle fiber CSA for each mouse using ImageJ.

#### Adipocyte CSA

Sections (4 µm) of iWAT and gWAT were cut and stained with hematoxylin and eosin to assess adipocyte size. Whole slides were scanned and 2-3 representative 500 µm x 500 µm images from each of two levels of the tissue were used to quantify the mean adipocyte CSA for each mouse using ImageJ.

#### RNA isolation and qPCR

Soleus and gastrocnemius muscles, liver, iWAT and gWAT depots, and lung tumor (*G12D*) or lung tissue organoids (WT;1 well of a 6-well plate) were used for RNA isolation. For muscle and adipose tissues, RNA extraction was conducted after tissues were mechanically homogenized (Polytron PT2100, Kinematica, Bohemia, NY, USA) in 1 mL of TRI reagent.

Thereafter, 200 μL of chloroform was added, the sample was vortexed, and centrifuged at 4°C. For adipose tissues, RNA extraction was completed using the RNeasy Lipid Tissue Mini Kit (Qiagen, Venlo, Netherlands), according to the manufacturer’s protocols, with the addition of the elective DNAse digestion steps. For muscle tissues, supernatants were transferred to a new tube and equal amounts of isopropanol and high salt solution (1.2 M sodium chloride, 0.8 M sodium citrate) were added to each sample. For soleus muscle, 0.5 μL of glycogen (ThermoFisher Scientific, Waltham, MA, USA) was added to the RNA precipitation step to increase RNA yield. Samples were incubated and then centrifuged at 4°C. Pellets were washed in 95% ethanol and RNA was reconstituted in nuclease-free water for cDNA synthesis or RNA-seq.

For cell samples, media was aspirated, and cells were washed twice with Hanks’ Buffered Saline Solution (HBSS). TRI reagent was added to each well of a 12-well plate and incubated for 2 minutes at room temperature. Lysates were transferred to a 2-mL tube and chloroform (1:4 chloroform:TRI reagent) was added. Samples were shaken vigorously and centrifuged at 4°C. Supernatants were removed, and pellets were resuspended in 500 μL of 75% ethanol, vortexed briefly to wash, and centrifuged at 12,000 rcf for 5 minutes at 4°C. Samples were reconstituted in nuclease-free water for cDNA analysis.

For tissue and cell samples, RNA quality and quantity were assessed (NanoDrop; ThermoFisher Scientific, Waltham, MA, USA) and RNA was reverse transcribed using qScript cDNA Supermix. qPCR was performed according to a standard protocol using iTaq Universal SYBR Green Supermix and validated primer sequences (**Supplemental Table S1**). Target gene expression was normalized to the average of two housekeeping genes (adipose: *Ppia, B2m*; muscle: *Gapdh, Ppia*). Results are expressed as 2^-ΔΔCt^ relative to the average of the housekeeping genes for each respective tissue and all data expressed relative to WT ^85^.

For organoid RNA isolation for RNA-seq, media was collected from cultured organoids and the adherent organoids were rinsed with phosphate-buffered saline (PBS). Over ice, Cell Recovery Solution (Corning, Corning, NY, USA) was added to the cells, cultures were agitated by repeated pipetting and incubated at 4°C. Samples were transferred to microcentrifuge tubes and centrifuged at 4°C. Supernatant was removed and organoid pellet was washed in PBS and centrifugation step was repeated. RNA isolation was conducted using the Direct-zol™ RNA MiniPrep kit (Zymo Research Corp., Irvine, CA, USA), according to manufacturer’s instructions, with Trizol (Zymo Research Corp, Irvine, CA, USA) and column-based spin steps to remove the hydrogel matrix. Following RNA isolation, RNA Clean & Concentrator™ kit (Zymo Research Corp., Irvine, CA, USA) was used, according to the manufacturer’s instructions to concentrate RNA for library preparation. RNA quality and quantity was confirmed by Qubit (ThermoFisher Scientific, Waltham, MA, USA).

#### RNA-seq

RNA libraries from isolated organoids were prepared from 500 ng of total RNA using the Zymo-Seq RiboFree Total RNA Library Kit according to the manufacturer’s instructions. RNA-Seq libraries were sequenced on an Illumina NovaSeq to a sequencing depth of at least 30 million read pairs (150 bp paired-end sequencing) per sample. Data were analyzed using the Zymo Research RNA-Seq pipeline, which was adapted from nf-core/rnaseq pipeline v1.4.2 ^86^ and built using Nextflow ^87^. Briefly, quality control of raw reads was conducted using FastQC v0.11.9. Adapter and low-quality sequences were trimmed from raw reads using Trim Galore! V0.6.6. Trimmed reads were aligned to the C57BL6/J mouse reference genome using STAR v2.6.1d. BAM file filtering and indexing was carried out using SAMtools v1.9 ^88^. RNA-seq library quality control was implemented using RseQC v4.0.0 ^89^ and QualiMap v2.2.2-dev ^90^. Duplicate reads were marked using Picard tools v2.23.9 ^91^. Library complexity was estimated using Preseq v2.0.3 ^92^. Duplication rate quality control was performed using dupRadar v1.18.0^93^. Reads overlapping with exons were assigned to genes using featureCounts v2.0.1^94^. Classification of rRNA genes/exons and their reads were based on annotations and RepeatMasker rRNA tracks from UCSC genome browser when applicable. Differential gene expression analysis was completed using DESeq2 v1.28.0 ^95^. Functional enrichment analysis were performed using g:Profiler python API v1.0.0 ^96^, the Broad’s GSEA software ^97^ and the R package PathfindR ^98^. Quality control and analysis results plots were visualized using MultiQC v1.9.

To confirm that organoid cultures maintained the Kras^G12D^ mutation, BAM files generated from organoid RNA-seq analyses were indexed with SAMtools v1.9 ^88^. The presence and frequency of Kras allelic variants were then visualized and quantified using the Integrated Genome Viewer (IGV) ^99^. Differentially expressed genes (DEGs) corresponding to WT and G12D lung organoid RNA-seq libraries were identified from raw read counts using DESeq2 and plotted with ggplot2. Finally, DEGs identified with DESeq2 were filtered for p-values adjusted for false discovery rate of <0.05 and absolute value of fold change >2 were subsequently cross-referenced against DEGs identified in the TRACERx *study* ^55^ to identify overlaps.

For analysis of adipose tissues, total RNA was isolated from gWAT and BAT from WT and G12D tissue samples collected at 3 and 6 weeks post-tamoxifen injection. RNA libraries were prepared using NEBNext Ultra II RNA Library Prep Kit for Illumina, following the manufacturer’s protocol. Libraries were sequenced on an Illumina NovaSeq platform at a depth of 30 million reads per sample (150 bp, paired-end). Resulting reads were aligned to the GRCm39 reference genome using STAR (v2.7.3.a) and, and gene-level quantification was performed with featureCounts (v2.0.0). Differential expression analysis was conducted using DESeq2 (v1.48.1), and gene set enrichment analysis (GSEA) was carried out using the FGSEA R package (v1.34.0).

#### Glycerol levels

Glycerol levels of mouse serum, human plasma, 3T3L1 adipocyte supernatants and lung tissue/tumor organoid culture conditioned media were measured using the Glycerol Assay Kit according to the manufacturer’s instructions and were expressed relative to body weight to control for CC-related wasting.

#### Hepatic triglycerides

Liver tissue (350-400 mg) was minced in substrate assay reagent. Hepatic triglyceride content was determined according to the kit manufacturer’s instructions and standardized to the starting tissue weight of the sample.

#### Serum metabolomics

Serum samples were thawed on ice and metabolites were extracted using ice-cold 5:3:2 methanol:acetonitrile:water ^100^. Mixtures were vortexed for 30 min at 4**°**C and centrifuged at 4**°**C. Samples were randomized then 10 μL of extracts were injected into a Thermo Vanquish UHPLC system (San Jose, CA, USA) and resolved on a Kinetex C18 column (150 × 2.1 mm, 1.7 μm, Phenomenex, Torrance, CA, USA) at 450 μL/min through a 5 min gradient from 0 to 100% B (mobile phases: A = 95% water, 5% acetonitrile, 1 mM ammonium acetate; B = 95% acetonitrile, 5% water, 1 mM ammonium acetate) in negative ion mode ^101^. Solvent gradient: 0-0.5 min 0% B, 0.5-1.1 min 0-100% B, 1.1-2.75 min hold at 100% B, 2.75-3 min 100-0% B, 3-5 min hold at 0% B. Injections were then repeated for positive ion mode at 450 μL/min through a 5 min gradient from 5 to 95% B (mobile phases: A = water, 0.1% formic acid; B = acetonitrile, 0.1% formic acid) in positive ion mode. Solvent gradient: 0-0.5 min 5% B, 0.5-1.1 min 5-95% B, 1.1-2.75 min hold at 95% B, 2.75-3 min 95-5% B, 3-5 min hold at 5% B. Eluant was introduced to the mass spectrometer (Thermo Q Exactive) using electrospray ionization. For both negative and positive polarities, signals were recorded at a resolution of 70,000 over a scan range of 65-900 m/z. The maximum injection time was 200 ms, microscans 2, automatic gain control (AGC) 3 x 10^6 ions, source voltage 4.0 kV, capillary temperature 320**°**C, and sheath gas 45, auxiliary gas 15, and sweep gas 0 (all nitrogen). Resulting .raw files were converted to .mzXML format using RawConverter. Metabolites were assigned and peak areas integrated using Maven (Princeton University) in conjunction with the KEGG database and an in-house standard library of >600 compounds ^100, 102^. Top metabolites were determined by t-test p-value, and data were normalized to median and auto-scaled in MetaboAnalyst.

#### UCP-1 detection in paraffin-embedded white adipose tissue

Paraffin-embedded sections were heated (70°C), deparaffinized in xylenes, washed in 100% ethanol, 95% ethanol, and PBS, and endogenous peroxidases quenched with 10% hydrogen peroxide in methanol. After washing in PBS, epitope retrieval was conducted at 95°C in citrate EDTA buffer (10 mM citric acid, 2 mM EDTA, 0.05% Tween 20, pH 6.2). Slides were rinsed in distilled water and PBS for 5 minutes and then blocked in 10% normal goat serum (Vector Laboratories S-1000-20) in PBS. Slides were incubated in primary antibody (1:1000 dilution in PBS) for 1 h at room temperature, rinsed in PBS, and secondary antibody (1:2000 dilution in PBS) was added. After washing in PBS, 3,3’-Diaminobenzidine (DAB) chromogen was added, slides were incubated in the dark and then rinsed in tap water. Slides were counter-stained in hematoxylin for 20 seconds and rinsed again in tap water until clear. Slides were dehydrated in serial ethanol solutions, mounted, and cover slipped. Slides were scanned at 40x magnification on a Leica-Aperio VERSA whole slide imager (Leica Biosystems, Wetzlar, Germany).

Images were analyzed using Halo software. The hue of the brown chromogen stain indicating positive UCP1 was segmented with the Halo module “Area Quantification version 2.2.1”. Stain selection was set to 2, stain 1 (hematoxylin) with stain color 0.239, 0.230, 0.144; stain 2 (UCP1) with stain color 0.139, 0.251, 0.216. Settings for stain detection were as follows: Hematoxylin markup color (255, 0, 0), hematoxylin Min OD (0.15, 0.15, 0.15), UCP1 markup color (0, 255, 0), UCP1 Min OD (0.15, 0.15, 0.15). Advanced settings were utilized to account for the negative space in the images (Min Tissue OD 0.037). Nuclei were counted using Halo module “Nuclei Detection – ISH IHC v 3.1.4.” General stain settings were utilized to detect the nuclear stain (hematoxylin – 0.438, 0.342, 0.171) and exclusion stain settings (0.110, 0.219, 0.189) were used to guard against non-specifically counting tissue lifting defects as nuclei. Typical settings on our samples were: Nuclear Contrast Threshold (0.497), Minimum Nuclear Optical Density (0.07), and Nuclear Size (1,1000). Default settings were used for Membrane and Cytoplasm Detection, and ISH cell scoring. Exclusion Marker threshold was set to 2. The Minimum Tissue OD setting was set to 0.003 to account for negative space in the images.

#### Statistics

Data were assessed for normality prior to statistical analyses. Graphical data are represented as the mean ± the standard error of the mean (SEM). ELISA data, if non-normally distributed, were log_10_ transformed prior to statistical analyses and the assumption of normality confirmed. Differences between WT and *G12D* groups for singular time point measurements were determined with a Student’s T-test, whereas differences over time were determined by repeated measures analysis of variance (ANOVA) with individual comparisons at each timepoint if group X time interaction effects were noted. For comparison of WT, *G12D,* and *KS* groups, a one-way ANOVA with multiple comparisons was used to determine differences between groups. Differences between these three groups over time were distinguished using a repeated measures one-way ANOVA with group X time interaction effects. *SPSS* was used for these analyses. Data and statistical analyses were visualized using Graphpad Prism.

For human NLST data analyses, we tabulated baseline summary statistics according to key demographics and study measurements for the exposed (lung cancer patients) and unexposed (lung cancer-free) arms of the NLST cohort. We visualized crude time trends in body composition measurements using both box plots and distributional dot plots overlaid with local regression trends. We then fit crude and multivariable hierarchical linear models to compare changes in the sum of skeletal muscle (SM), subcutaneous adipose tissue (SAT) and visceral adipose tissue (VAT) area between lung cancer cases and matched non-cases across the fifth, eight, and tenth thoracic vertebral levels, indexed to patient height and tissue attenuation and stratified by sex. All outcomes were regressed on the 3-way interaction between exposure (lung cancer/stage vs. non-case), measurement time (continuous), and participant sex; multivariable models further adjusted for race and ethnicity (dichotomized as non-Latino White vs. all other races and ethnicities due to sparse data), body mass index (standard categories modeled as a factor variable), and age (modeled as a continuous variable). All models included study center and patient ID as random effects to account for clustering by study site and correlation of within-patient measurements, respectively. We used changes in Akaike’s information criterion (AIC) to guide model specification. We specified contrasts to estimate p-values for overall differences in body composition slopes by exposure group within sex strata. Modeled data were visualized by plotting marginal predicted values across the range of time values by exposure group and stratified by sex, with standard errors estimated by the delta method. All analyses were performed with Stata v.18.5 (Stata Corporation, College Station, TX).

## Additional resources

None.

