## Supplemental Figures for "Early adipose tissue wasting in a novel preclinical model of human lung cancer cachexia"

**Supplementary Materials**

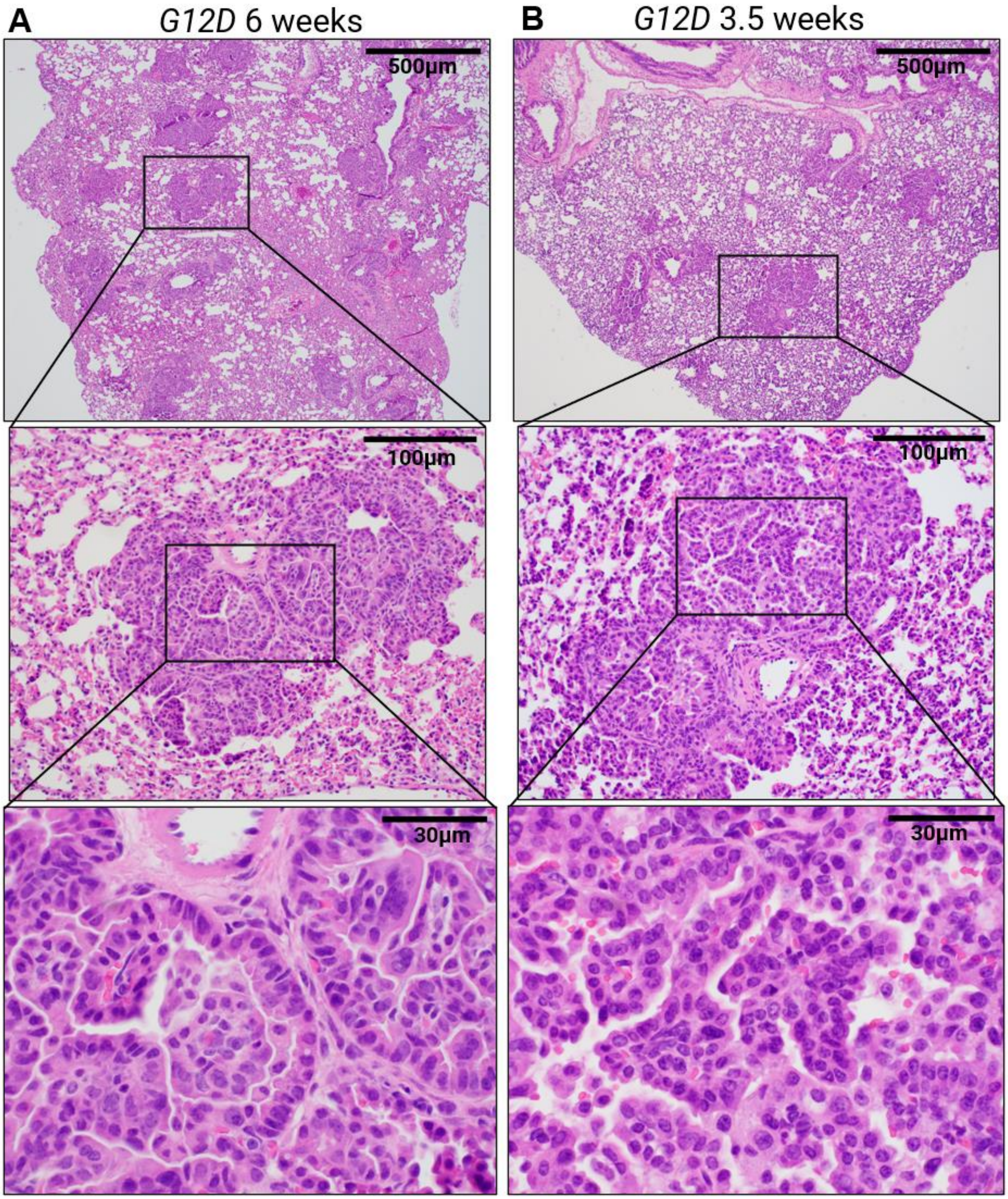

**Supplemental Figure S1. Evidence for adenocarcinomas in teminal bronchioles of lung tissue of *Kras*$^{G12D/+}$ (*G12D*) mice at 6- and 3.5-weeks post-induction, related to Figure 1.** Haematoxylin and eosin-stained images of lung tissue at 6 (A) and 3.5 weeks (B) post tamoxifen showing adenocarcinoma development in the terminal bronchioles.

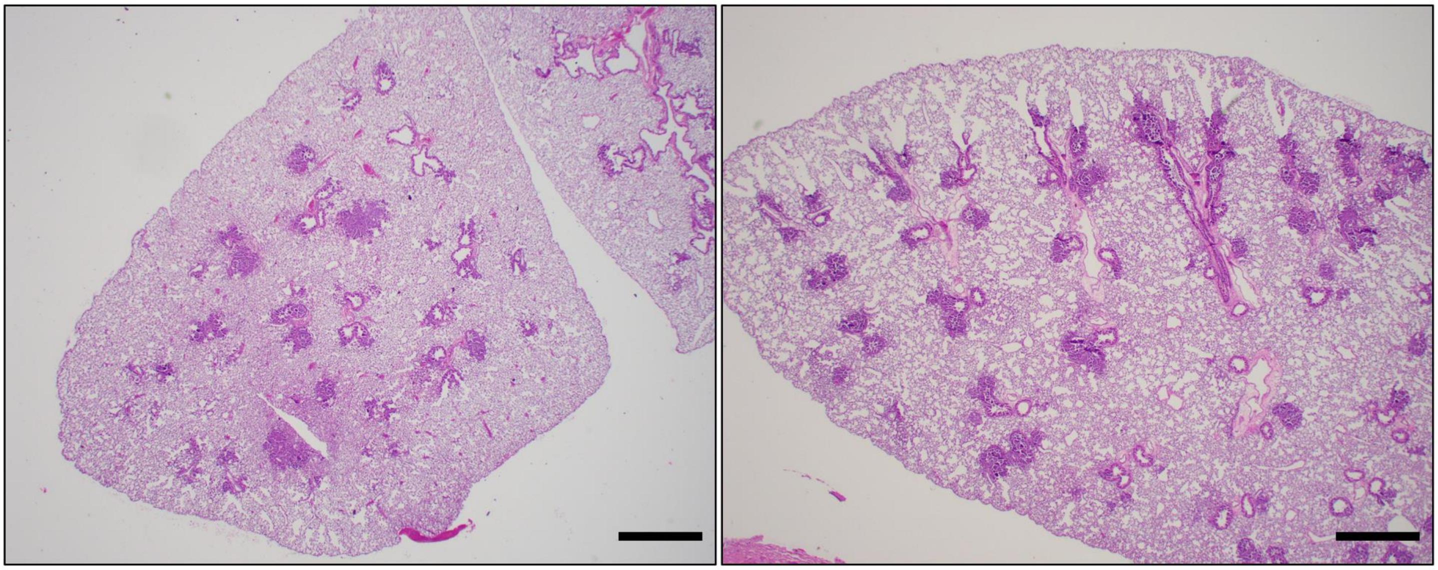

**Supplemental Figure S2. Non-tamoxifen-inducible (*i.e.,* leaky) Cre expression causes eventual lesion development in Kras$^{G12D/+}$ (*G12D*) mice at 25 weeks of age, related to Figure 1**. Haematoxylin and eosin-stained lungs of 25-week-old *G12D* mice in the absence of tamoxifen administration. Scale bars: 500 µm.

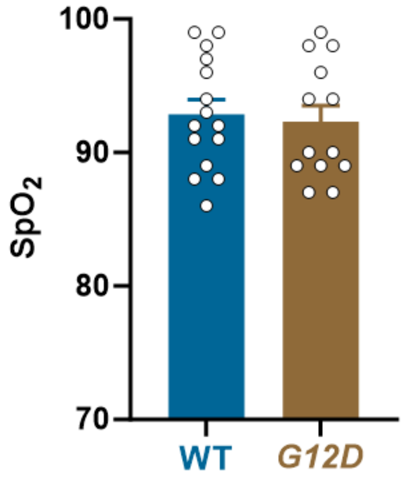

**Supplemental Figure S3. Effect of lung epithelial cell specific *Kras*$^{G12D/+}$ (*G12D*) on blood oxygen saturation ($SpO_2$), related to Figure 1.** $SpO_2$ was measured by pulse oximetry in wild type (WT; blue bar) and *G12D* (gold bar) mice 3 weeks post-induction. Data were analyzed by Student's t-test.

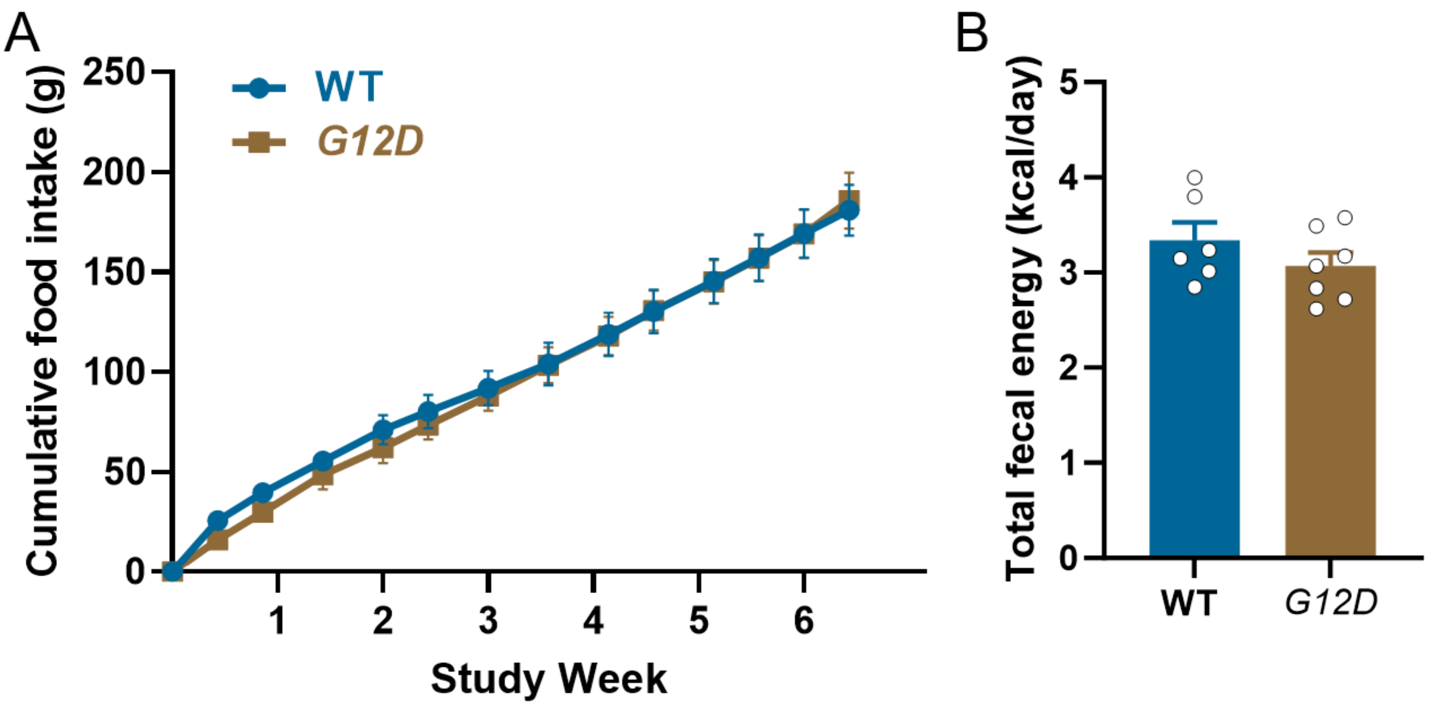

**Supplemental Figure S4. Effect of lung epithelial cell specific *Kras*$^{G12D/+}$ (*G12D*) on food intake prior to initiation of weight loss, related to Figure 1.** In a cohort of singly housed wild type (WT; n=3; blue circles/line) and *Kras*$^{G12D}$ (*G12D*) mice (n=4-5; gold squares/line), we monitored food intake and body weight for 6 weeks post-induction to examine changes in food consumption prior to the divergence of body weight (see **Fig. 1C**). Food intake was measured twice weekly across the 6-week period and is displayed as cumulative intake (A), analyzed using a repeated measure analysis of variance. Fecal energy content from a 72-hour period collected during week 6 was measured by bomb calorimetry in separate cohorts of *G12D* (n=7) and WT (n=6) mice (B), analyzed by Student's T-Test.

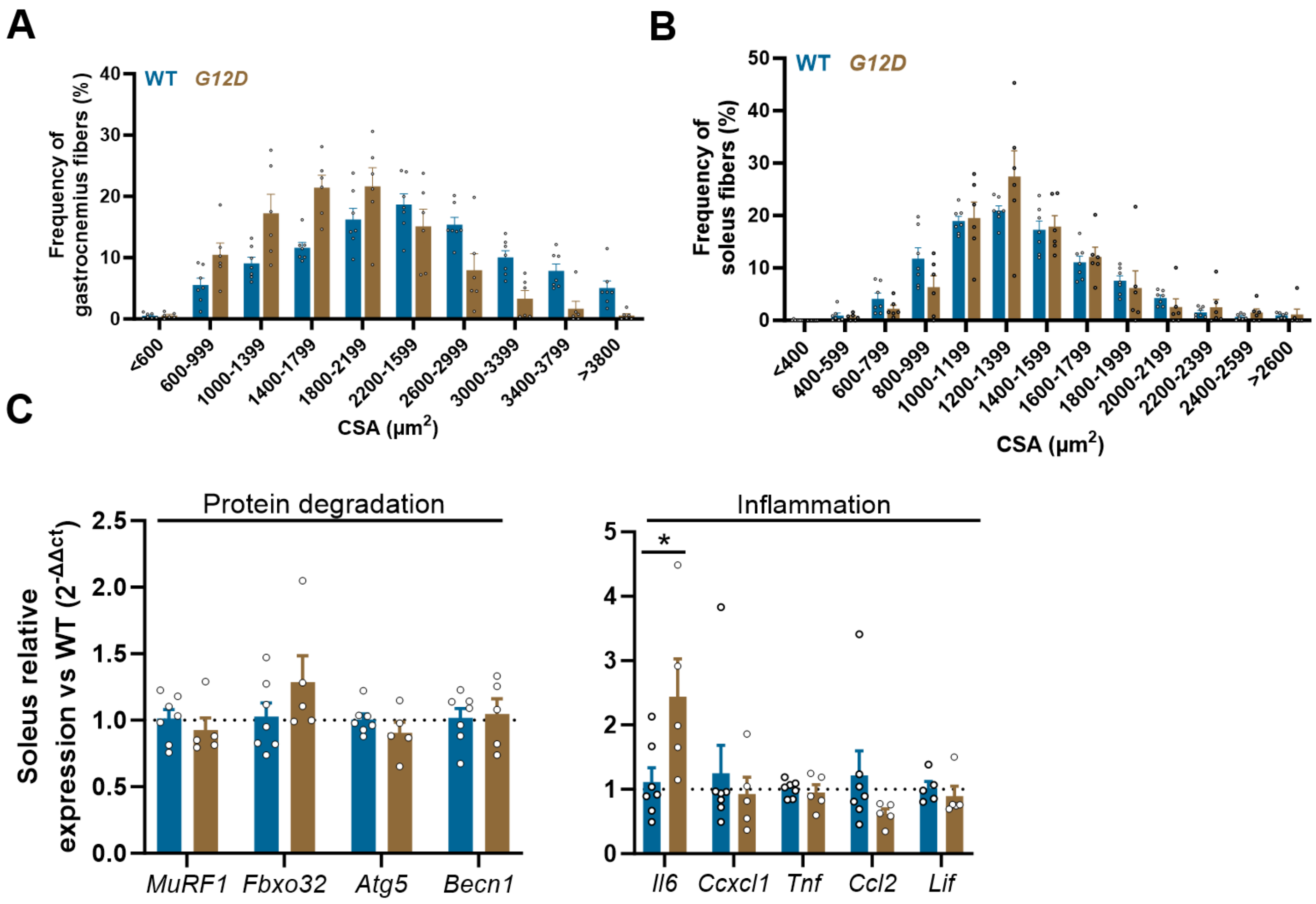

**Supplemental Figure S5. Effect of lung epithelial cell specific *Kras*$^{G12D/+}$ (*G12D*) on gastrocnemius and soleus muscle cross-sectional area and soleus mRNA expression levels at 12-weeks post-induction, related to Figure 2.** Frequency (%) distribution of gastrocnemius (A) and soleus (B) fiber cross-sectional area in wild type (WT; n=7; blue bars) and *G12D* mice (n=6; gold bars) 12 weeks post-induction for soleus muscle. Gene expression of skeletal muscle markers of proteolytic degradation and inflammation in soleus muscle (C). Data were analyzed by Student's t-test, with * $p$<0.05.

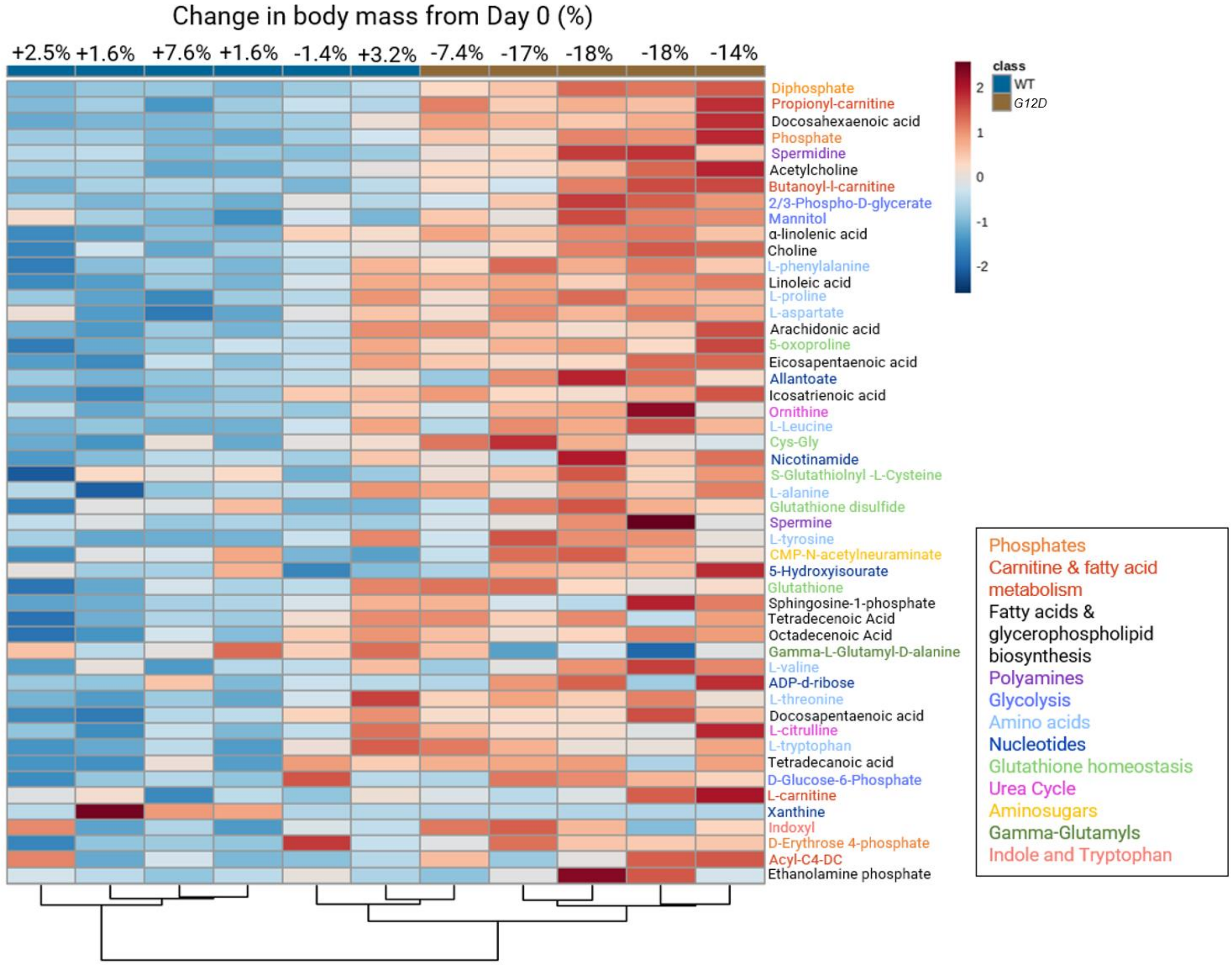

**Supplemental Figure S6. Effect of lung epithelial cell specific *Kras*$^{G12D/+}$ (*G12D*) on serum metabolites 12 weeks post-injection, related to Figure 2.** Hierarchical clustering analysis of the top 50 serum metabolites by *p*-value (mean of each group compared by t-test, with $p<0.05$) between *G12D* (n=5; gold bars at top of columns) and wild-type littermates (WT; n=6; blue bars at top of column) after 12 weeks of tumor development with the relative change in body weight over the 12-week study period listed above each animal. Data were normalized to median and auto scaled in MetaboAnalyst.

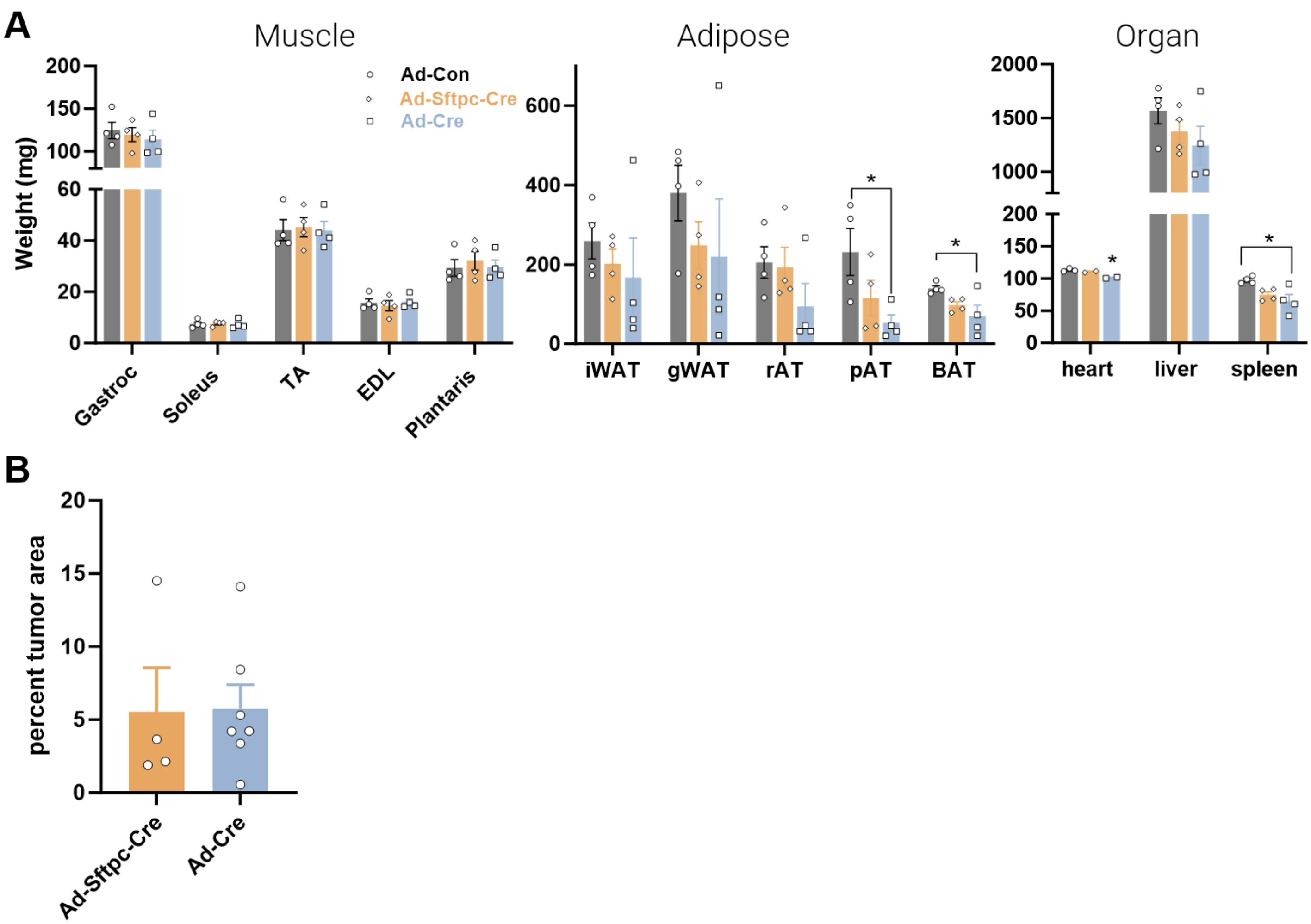

**Supplemental Figure S7. Tissue and organ weights in mice with *Kras*$^{G12D}$ induced by oropharyngeal administration of adenovirus specific to AT2 cells or one that induces the transgene across lung epithelial cell types at 12 weeks post-injection, related to Figure 2**. Cre recombinase was introduced to induce tumor development in *Kras*$^{G12D/+}$ mice using AT-2 cell-targeted, adenovirus (Ad-Sftpc-Cre; peach bars/open diamonds) or adenovirus Ad5-CMV-Cre targeting all airway epithelial cells (Ad-Cre; light blue bars/open squares), and were compared with control adenovirus delivery (Ad-Con; grey bars/open circles) that did not develop lung tumors. Tissues were collected (12-16 weeks post Ad instillation) to determine muscle, fat pad, and organ weights (A) and quantify tumor area in sectioned lung (B).

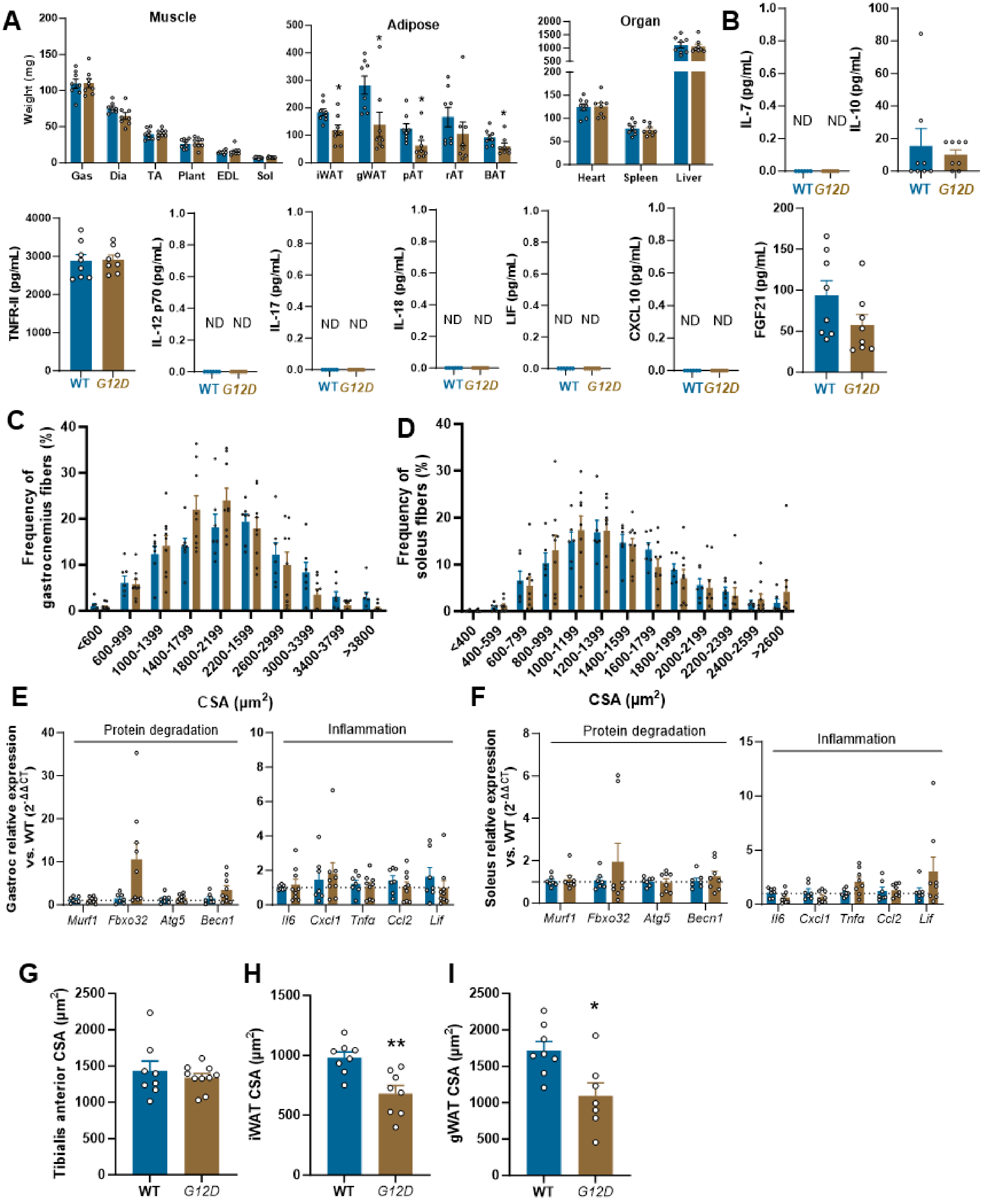

**Supplemental Figure S8. Effect of lung epithelial cell specific *Kras*$^{G12D/+}$ (*G12D*) on muscle and fat weight, additional serum cytokines, muscle fiber size and distribution and gene expression and fat cell**

**size 6 weeks post-induction, related to Figure 4.** In a separate cohort of wild-type (WT; n=8; blue bars) and *G12D* (n=8; gold bars) mice (same mice used in Fig. S4B), we measured the weight of additional hindlimb muscles and adipose depots 6 weeks post-injection (A) and additional circulating inflammatory mediators (B). Frequency distribution of gastrocnemius (C) and soleus (D) muscle fiber cross-sectional areas (CSA; scale bars=100 µm) in the mice shown in Fig. 4: WT (n=7; blue bars) and *G12D* (n=8; gold bars). Gene expression in gastrocnemius (E) and soleus muscle (F) in mice show in Fig.4. In mice from Fig S4B, muscle fiber size was measured in tibialis anterior muscle (G), and adipocyte size in iWAT (H) and gWAT (I). Data were analyzed by Student's t-test with $*P<0.05$ ; ND=not detected. IL-7: interleukin-7; IL-10: interleukin-10; TNFRII: TNF receptor II; IL-12: interleukin-12; IL-17: interleukin-17; IL-18: interleukin-18; LIF: leukocyte inhibitory factor; CXCL10: C-X-C motif chemokine ligand 10; FGF21: fibroblast growth factor 21.

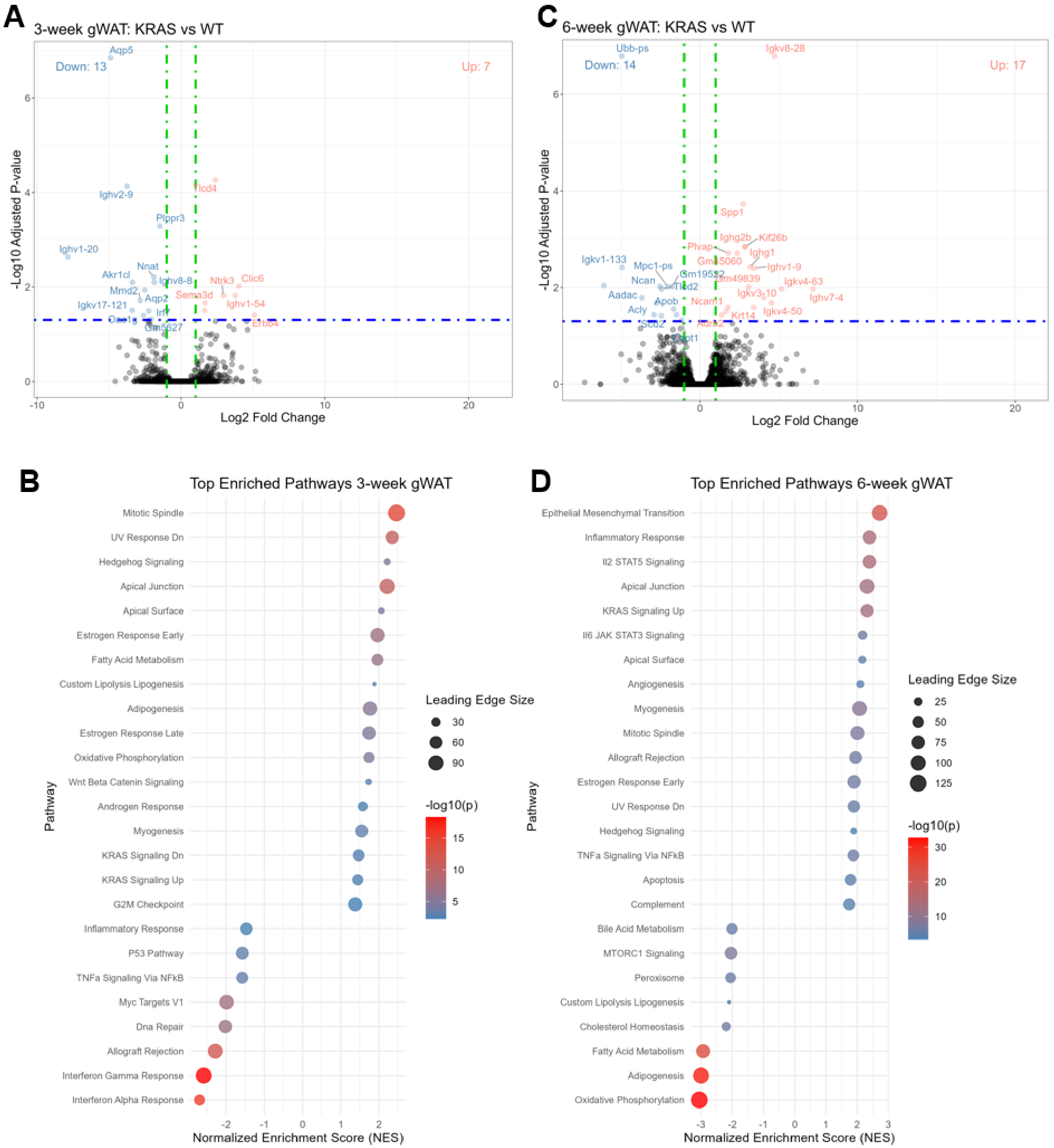

**Supplemental figure S9. Transcriptional profiling of perigonadal white adipose tissues (gWAT) of *Kras*$^{G12D/+}$ (*G12D*) mice at 3 and 6 weeks post-induction, related to Figure 4.** gWAT adipose tissue RNA was isolated and *RNAseq* was carried out to determine transcriptional changes. Volcano plot depicting differentially expressed genes, with red representing upregulated (n=7) and blue representing downregulated genes (n=13) in *G12D* gWAT compared to WT gWAT at 3 weeks post-injection, with a *p*-value exclusion of 0.05 (A). Pathway analysis using Gene Set Enrichment Analysis (GSEA) depicting differentially expressed pathways in *G12D* gWAT in comparison with WT gWAT at 3 weeks post-injection (B). Volcano plot depicting differentially upregulated (n=17) and downregulated (n=14) genes in *G12D* gWAT compared to WT gWAT at 6 weeks post-injection, with a *p*-value exclusion of 0.05 (C). Pathway analysis using GSEA depicting differentially expressed pathways in *G12D* gWAT in comparison with WT gWAT at 6 weeks post-injection (D). A custom pathway list was created for '*Lipolysis Lipogenesis'*; the genes included in this list are in **Supplemental Table S1.**

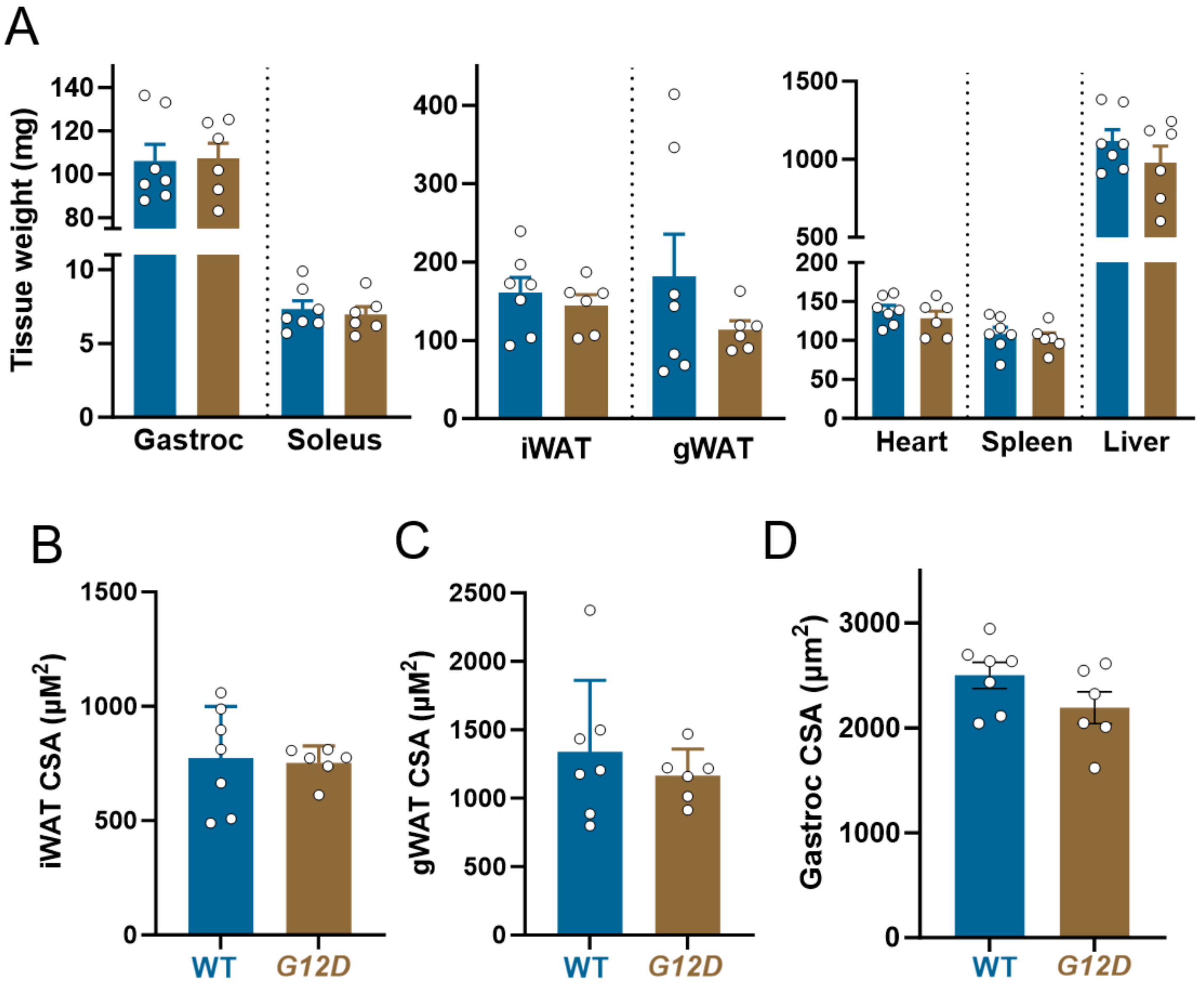

**Supplemental Figure S10. Effect of lung epithelial cell specific *Kras*$^{G12D/+}$ (*G12D*) on tissue weights and adipose and skeletal muscle cross-sectional area (CSA) 3 weeks post-induction, related to Figure 4.** Muscle (*Gastroc*, gastrocnemius), adipose tissues, and organ weights (A), inguinal white adipose tissue (iWAT; B) perigonadal (gWAT; C) and gastrocnemius (Gastroc) CSA (D) in wild type (WT; n=7; blue bars) and *G12D* (n=6; gold bars). Data were analyzed by Student's t-test.

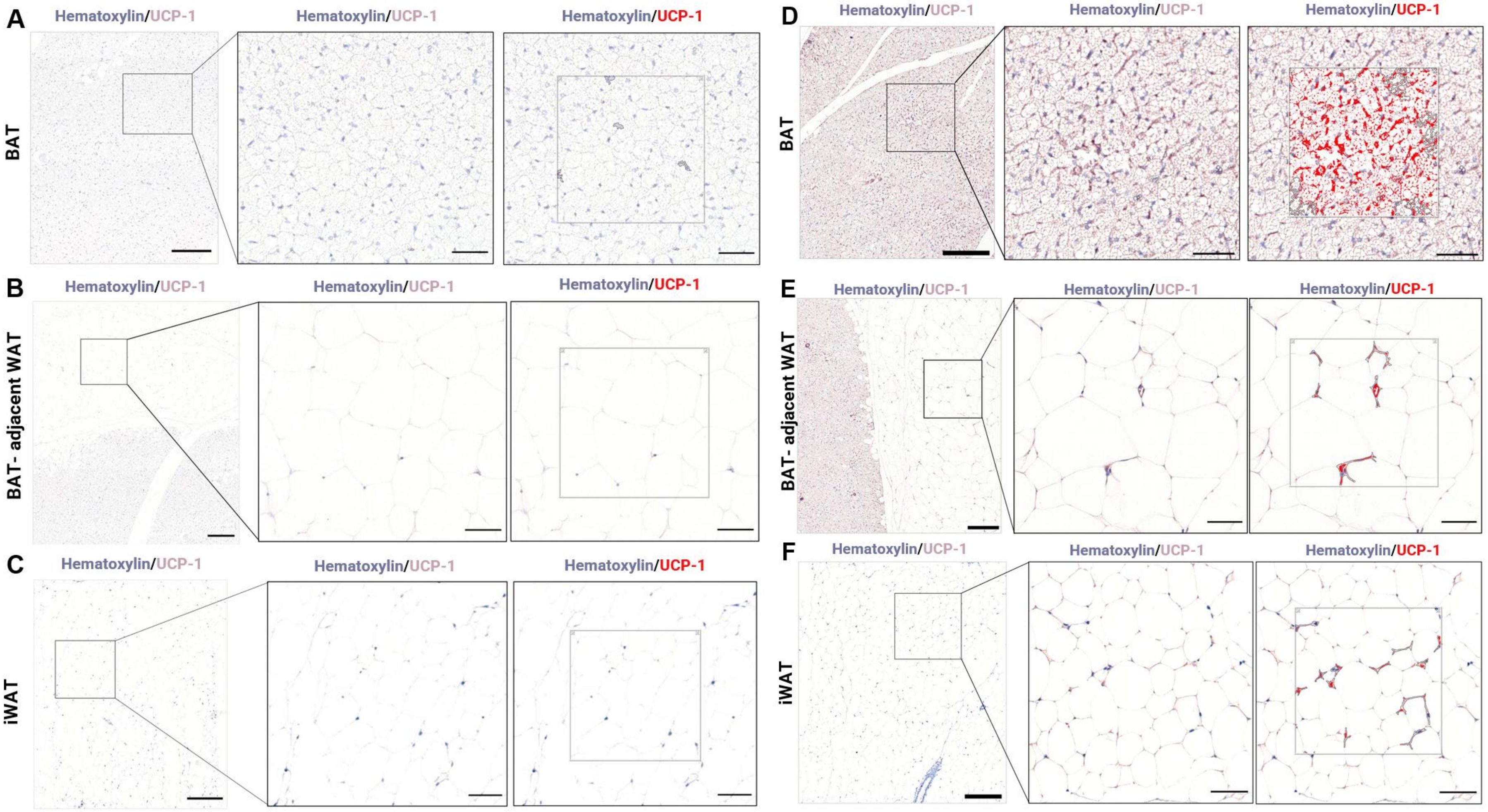
A
BAT
Hematoxylin/UCP-1
Hematoxylin/UCP-1
Hematoxylin/UCP-1
B
BAT- adjacent WAT
Hematoxylin/UCP-1
Hematoxylin/UCP-1
Hematoxylin/UCP-1
C
iWAT
Hematoxylin/UCP-1
Hematoxylin/UCP-1
Hematoxylin/UCP-1
D
BAT
Hematoxylin/UCP-1
Hematoxylin/UCP-1
Hematoxylin/UCP-1
E
BAT- adjacent WAT
Hematoxylin/UCP-1
Hematoxylin/UCP-1
Hematoxylin/UCP-1
F
iWAT
Hematoxylin/UCP-1
Hematoxylin/UCP-1
Hematoxylin/UCP-1

**Supplemental Figure S11. Immunohistochemical analysis of adipose tissue uncoupling protein 1 (UCP1) content, related to Figure 4.** Images depict signal detection for UCP1 (brown) in adipose tissue sections co-stained with hematoxylin (blue; left and center images of each panel) and rendered via HALO software for quantitation (red; right-most image). Images representing negative controls in the absence of primary antibody in interscapular brown adipose tissue (BAT; A), WAT adjacent to BAT (B) and iWAT (C). Representation of UCP-1 positive signal in interscapular BAT is shown as a positive control (D), in WAT adjacent to BAT (E), which was used to set signal detection threshold, and in iWAT (F). Enlarged images show greater detail of hematoxylin and UCP1 staining (central images) and UCP1 signal derived from image post-processing using the HALO software (right-most images). Scale bars in the left-most columns are 200 μM and those in the center and right-most columns are 50 μM.

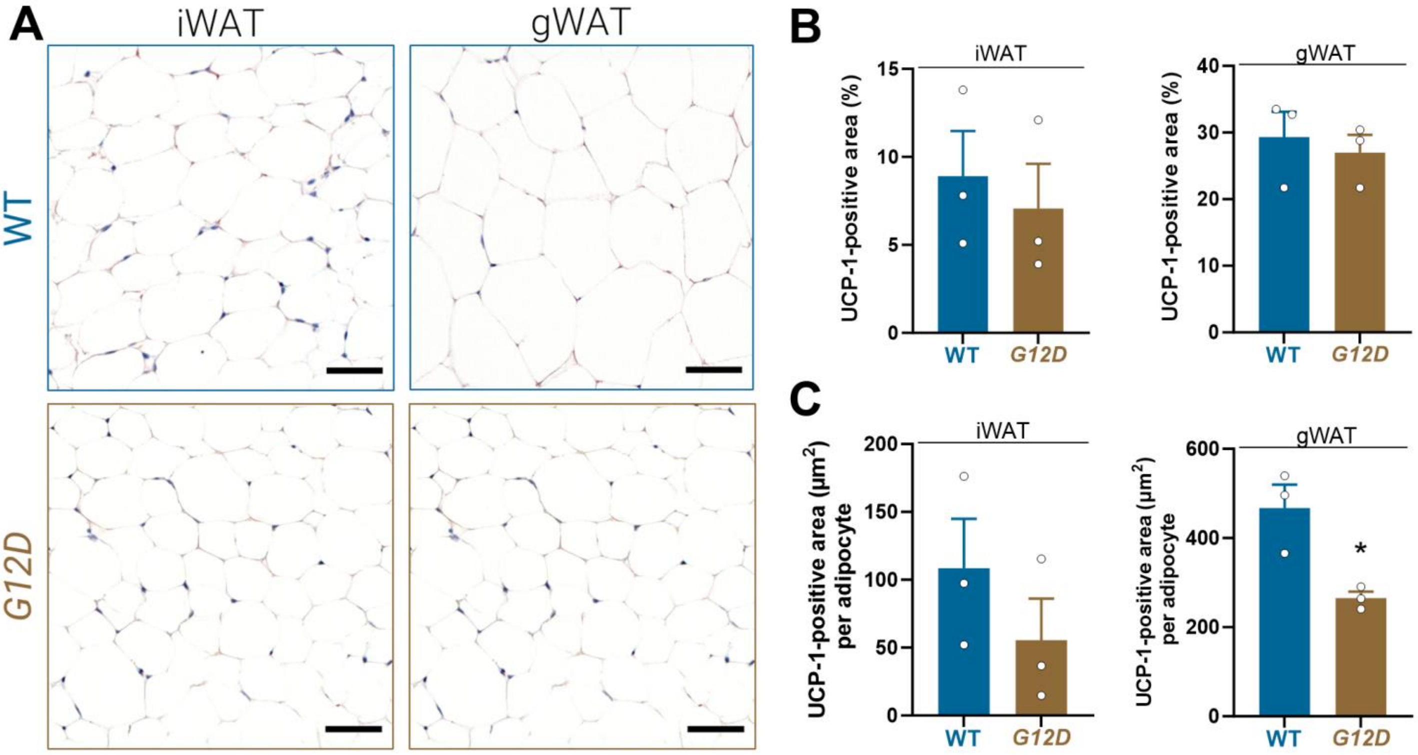

**Supplemental Figure S12. Effect of lung epithelial cell specific *Kras$^{G12D/+}$ (G12D)* on uncoupling protein-1 (UCP1) expression in adipose tissue depots during early CC development, related to Figure 4.** Representative images of immunohistochemistry against UCP1 in iWAT and gWAT tissues in 6-week WT (n=3; blue bars) and *G12D* mice (n=3; gold bars; scale bars = 50 μm) (A). UCP1 positive area data expressed as percent of the total area (B) and standardized to average adipocyte size (C). Data were analyzed using Student's t-tests, with * $p$<0.05.

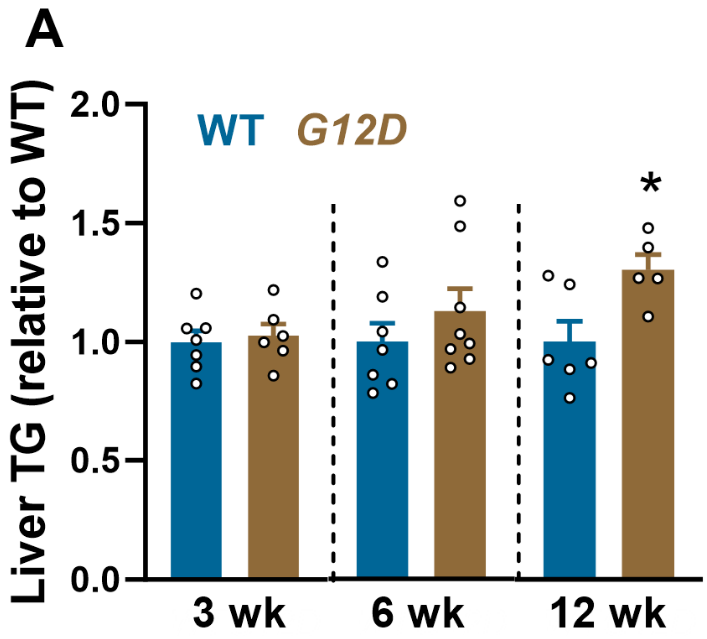

**Supplemental Figure S13. Effect of lung epithelial cell specific *Kras*$^{G12D/+}$ (*G12D*) on liver triglyceride (TG) levels at 3, 6 and 12 weeks post-induction, related to Figure 4.** Triglycerides (TG) were measured at 3, 6 and 12 weeks post tamoxifen-injection in wild-type (WT; blue bars, n=6-7/group) and *G12D* (gold bars; n=6-8 mice/group)(A). Data were analyzed using Student's t-tests, with *P<0.05.

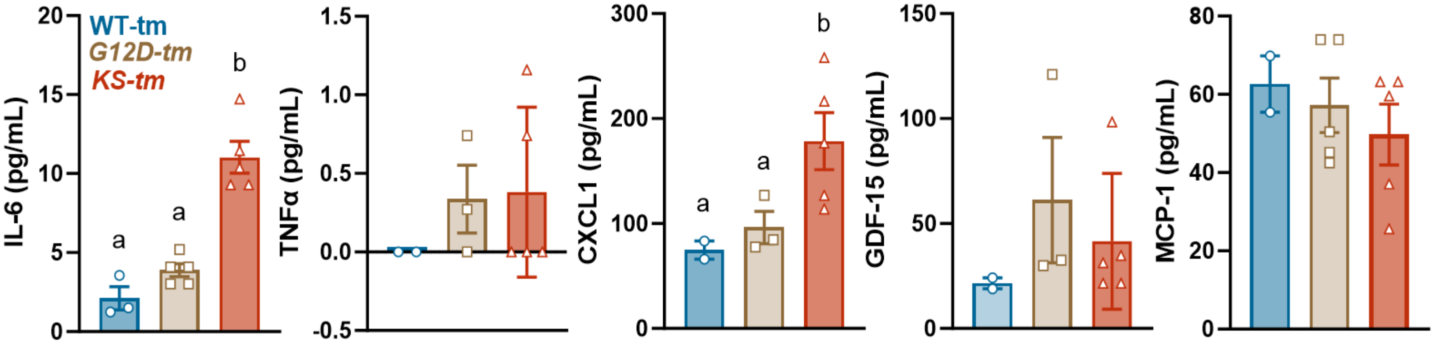

**Supplemental Figure S14. Comparison of serum cytokines in lung epithelial cell specific, inducible *Kras*$^{G12D}$ (*G12D-tm*) and *Kras*$^{G12D/+}$-*Stk11*$^{-/-}$ (*KS-tm*) murine models of CC, related to Figure 5.** Serum inflammatory cytokines of WT (n=4; blue bars/circles), *G12D-tm* (n=5; gold bars/squares) and *KS-tm* (n=5; red bars/triangles) mice injected with tamoxifen at 9-10 weeks of age and studied 24 days post-induction. Data were analyzed using one-way ANOVA with post hoc comparisons for those with $p<0.05$ group or group X time interaction effects. Different letters indicate differences by multiple comparisons P<0.05.

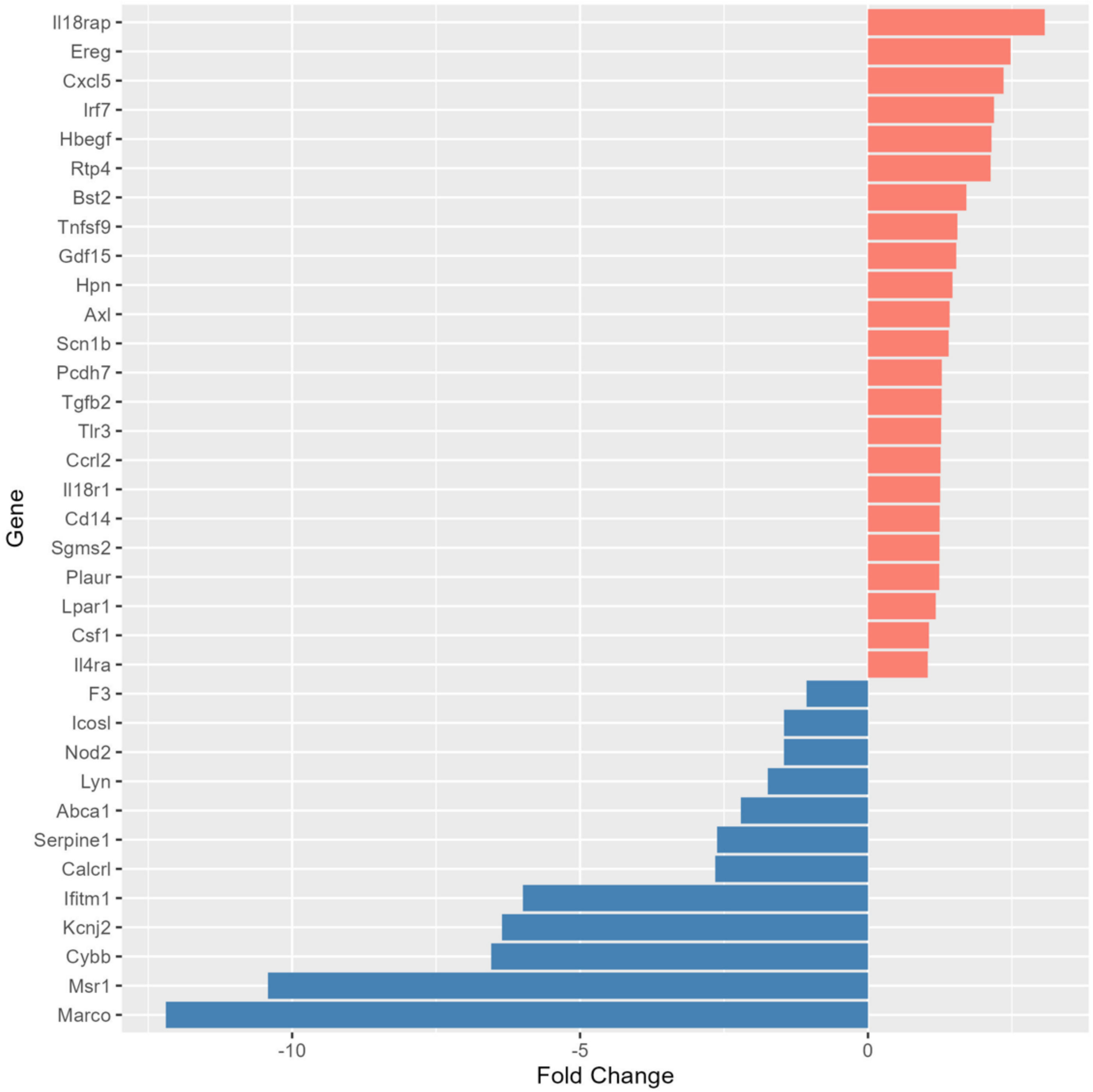

**Supplemental Figure S15. Exploration of transcriptional differences in inflammatory gene products associated with CC based on prior literature in organoids developed from lung tissue from wild-type (WT) and lung tumors from *Kras*$^{G12D/+}$ (*G12D*) mice, related to Figure 6.** Differentially expressed genes (DEGs) associated with inflammatory responses, with displayed genes filtered for *p*-values adjusted for false discovery rate of < 0.05 and absolute value of fold change >2.

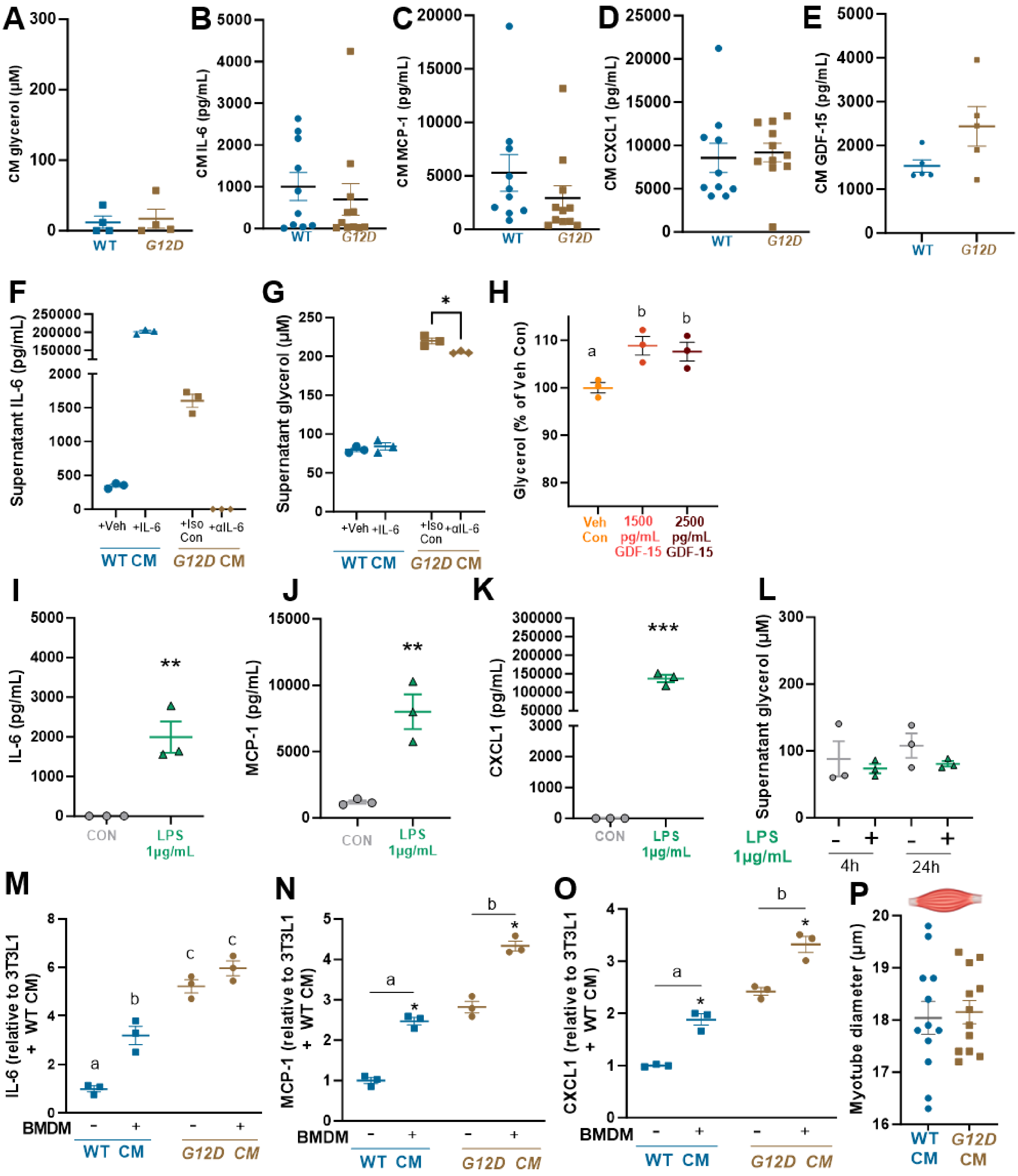

**Supplemental Figure S16. Supernatant glycerol and pro-inflammatory cytokines levels, in adipocytes treated with *G12D* lung tumor organoid conditioned media (CM), related to Figure 6.** CM glycerol (A), IL-6 (B), MCP-1 (C), CXCL1 (D) and GDF-15 (E) in samples used for CM treatment studies. 3T3L1 adipocytes were treated for 24 hours with WT CM with vehicle (+Veh) or recombinant mouse IL-6 (+IL-6) to determine the sufficiency of IL-6 to cause inflammation and lipolysis or with *G12D* CM treated with isotope control (+Iso Con) or neutralizing anti-IL-6 antibody (αIL-6) to determine the necessity of IL-6 to alter supernatant IL-6 (F) and glycerol (G). 3T3L1 adipocytes were treated with recombinant mouse GDF-15 at levels present in WT (1,500 pg/mL) or *G12D* (2,500 pg/mL) CM and supernatant glycerol measured after 24h (H). 3T3L1 adipocytes were treated with LPS (1 μg/mL; green triangles) or vehicle control (grey circles) for 24 hours and supernatant IL-6 (I), MCP-1(J), CXCL-1(K), and glycerol (L) levels were measured after 24 hours. Bone marrow-derived macrophages isolated from WT mice were co-cultured with 3T3L1 adipocytes and treated with WT or *G12D* CM. Supernatant IL-6 (M), MCP-1 (N), and CXCL1 (O) were measured after 24 hours of treatment. In differentiated C2C12 myotubes, myotube diameter was measured after 72 hours of WT or *G12D* CM treatment (1:3 dilution in differentiation medium). Analysis was Student's t-test (A-E, I-L, P), one-way ANOVA with multiple comparisons and letters indicating significant differences (L)  or two-way ANOVA (F-G, M-O), with post hoc comparisons for ANOVAs for those with $p<0.05$ group or group X BMDM interaction effects with different letters indicating differences and asterisks indicating a BMDM effect.

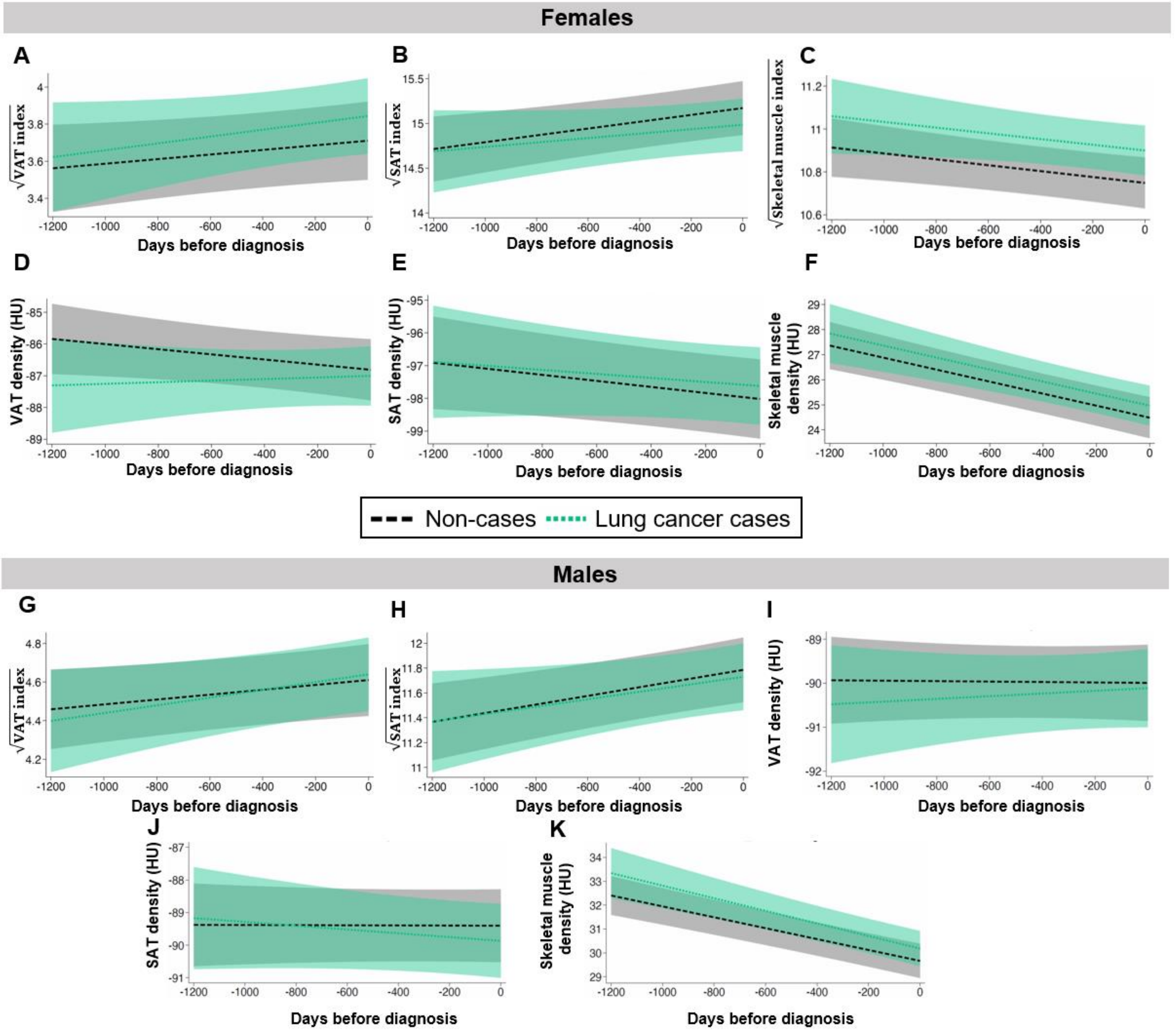

**Supplementary Figure S17. Multivariable hierarchical linear modeling of changes in visceral (VAT) and subcutaneous adipose tissue (SAT) and skeletal muscle among female (upper panels) and male (lower panels) non-case controls and lung cancer patients (green lines) prior to diagnosis in the National Lung Screening Trial, related to Figure 7.** Visceral (VAT) and subcutaneous adipose tissues (SAT) areas were indexed to patient height (Panels A-C, G,H) and density measured by tissue attenuation (Panels D-F, I-K; Hounsfield units; HU). Data were transformed via square root to meet the assumption of normality (Panels A-C, G-H). Data for VAT index,  SAT index, skeletal muscle index, VAT attenuation, SAT attenuation, and skeletal muscle attenuation are shown for females (A-F, respectively) are shown for non-case controls (black lines) and lung cancer cases (green lines). In Males, VAT index (G), SAT index (H), VAT attenuation (I), SAT attenuation (J), and skeletal muscle attenuation (K) are shown for non-case controls (black lines( and lung cancer cases (green lines). Modeled data were visualized by plotting marginal predicted values across the range of time values by exposure group. Standard errors were estimated by the delta method and 95% confidence intervals shown as the shaded regions of the corresponding color around each line. None of the analyses for these variables uncovered statistically significant differences.

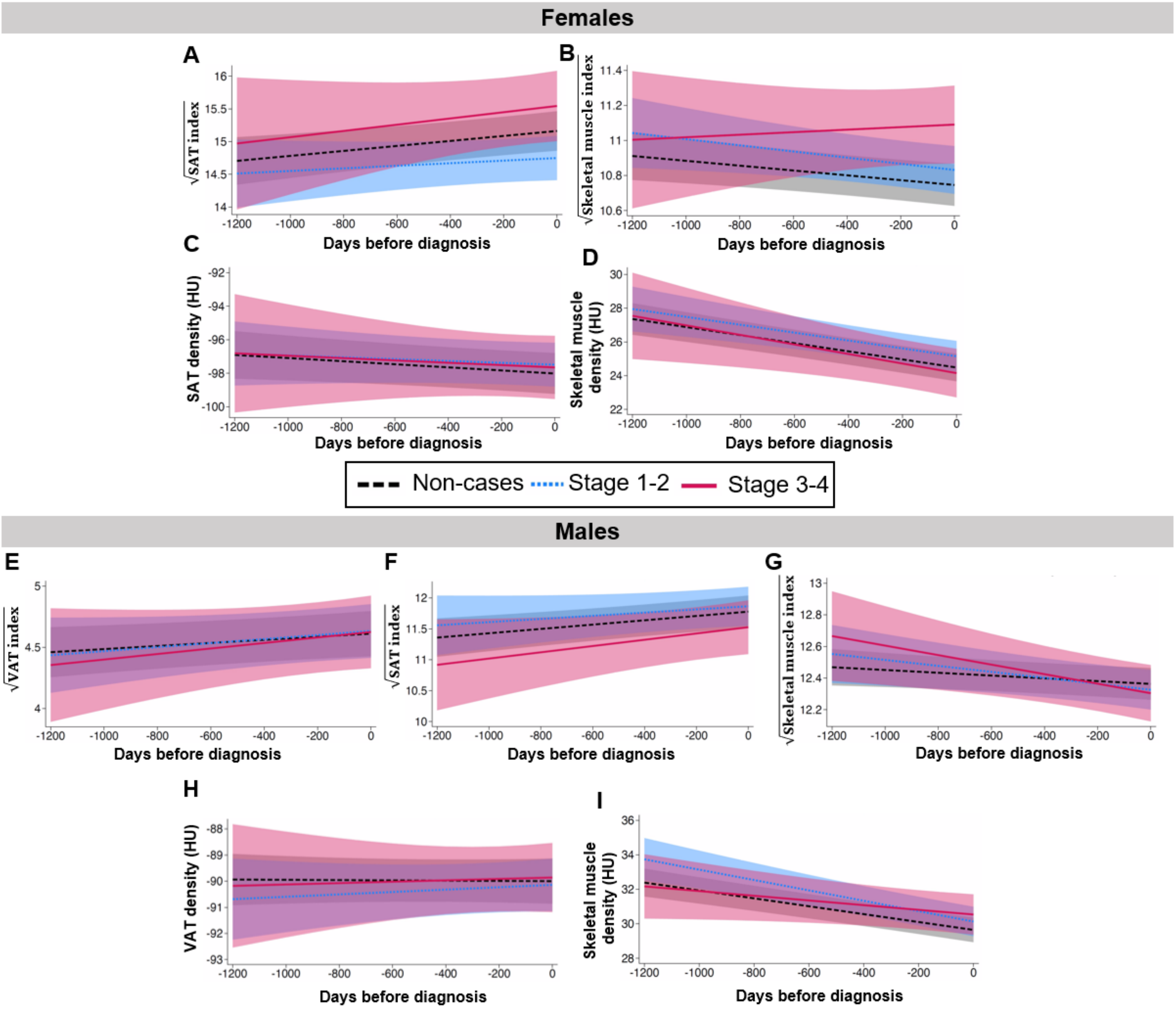

**Figure S18. Multivariable hierarchical linear modeling of changes in visceral (VAT) and subcutaneous adipose tissue (SAT) and skeletal muscle among female (upper panels) and male (lower panels) non-case controls and lung cancer patients stratified by cancer stage at diagnosis in the National Lung Screening Trial, related to Figure 7.** Visceral (VAT) and subcutaneous adipose tissues (SAT) areas were indexed to patient height and density measured by tissue attenuation (C,D, H-I; Hounsfield units; HU). Data were transformed via square root to meet the assumption of normality (A,B, E-G). Patients with lung cancer were stratified based on stage, with early-stage (1-2; blue lines) and advanced-stage (3-4; pink lines) compared to non-case controls (black lines). Modeled data were visualized by plotting marginal predicted values across the range of time values by exposure group. Standard errors were estimated by the delta method and 95% confidence intervals shown as the shaded regions of the corresponding color around each line. None of the analyses for these variables uncovered statistically significant differences.

**Supplemental Table S1.** Custom pathways employed in Gene Set Enrichment Analysis or RNAseq data from perigonadal white adipose tissues at 3 and 6 weeks post- *Kras*$^{G12D}$ induction

| **Custom Gene Set Name** | **Gene name** | **Protein product** |
|---|---|---|
| Creatine Metabolism | *Ppp1r1a* | Protein phosphatase 1 regulatory inhibitor subunit 1a |
| | *Kyat4* | Glutamic-oxaloacetic transaminase |
| | *Tead4* | TEA domain transcription factor 4 |
| | *Hoxa9* | Homeobox a9 |
| | *Anxa8* | Annexin A8 |
| | *Gatm* | Glycine amidinotransferase |
| | *Slc6a8* | Solute carrier family 6 member 8 |
| | *Gamt* | Guanidinoacetate N-methyltransferase |
| | *Ckmtl* | Citrate synthase lysine methyltransferase |
| | *Ckmt2* | Creatine kinase, mitochondrial 2 |
| | *Ckb* | Creatine kinase b |
| Calcium Cycling | *Atp2a2* | ATPase sarcoplasmic/endoplasmic reticulum Ca2+ transporting 2 |
| | *Atp2a3* | ATPase sarcoplasmic/endoplasmic reticulum Ca2+ transporting 3 |
| | *Ryr2* | Ryanodine receptor 2 |
| | *Pln* | Phospholamban |
| | *Arln* | Allregulin |
| | *Itpr3* | Inositol 1,4,5-triphosphate receptor 3 |
| | *Adra1a* | Adrenoceptor alpha 1A |
| | *Adora3* | Adenosine A3 receptor |
| | *Casq1* | Calsequestrin 1 |
| Lipogenesis / Lipolysis | *Cidea* | Cell death inducing DFFA like effector A |
| | *Pnpla2* | Patatin like domain t, triacylglycerol lipase (Adipose triglyceride lipase) |
| | *Lipe* | Hormone sensitive lipase |
| | *Acc* | Acetyl coA carboxylase |
| | *Prkaa1* | Protein kinase AMP-activated catalytic subunit alpha 1 |
| | *Prkaa2* | Protein kinase AMP-activated catalytic subunit alpha 2 |
| | *Prkab1* | Protein kinase AMP-activated catalytic subunit beta 1 |
| | *Prkab2* | Protein kinase AMP-activated catalytic subunit beta 2 |
| | *Prkag1* | Protein kinase AMP-activated catalytic subunit gamma 1 |
| | *Prkag3* | Protein kinase AMP-activated catalytic subunit gamma 2 |
| | *Acly* | ATP citrate lyase |
| | *Fasn* | Fatty acid synthase |
| | *Gpam* | Glycerol-3-phosphate acyltransferase, mitochondrial |
| | *Hadh* | Hydroxyacyl-coA dehydrogenase |
| | *Pcx* | Pyruvate carboxylase |
| | *Scd1* | Stearoyl-coenzyme A desaturase 1 |
| | *Aspg* | Asparaginase |
| | *Hsd17b10* | Hydroxysteroid 17-beta dehydrogenase 10 |
| | *Apoc3* | Apolipoprotein C3 |
| | *Acot1* | Acyl-CoA thioesterase 1 |
| | *Acot5* | Acyl-CoA thioesterase 5 |
| | *Mccc2* | Methylcrotonoyl-Coenzyme A carboxylase 2 |

**Supplemental Table S2.** Baseline characteristics of lung cancer cases and matched non-cases.

| Characteristic | Lung cancer cases (n=436) | | Non-cases (n=436) | |
|---|---|---|---|---|
| Age (median, IQR) | 64 | (59, 68) | 60 | (57, 65) |
| Age category, n (%) | | | | |
| <60 | 113 | (26) | 197 | (45) |
| 60-64 | 138 | (32) | 119 | (27) |
| 65-69 | 112 | (26) | 87 | (20) |
| ≥70 | 73 | (17) | 33 | (7.6) |
| Sex, n (%) | | | | |
| Male | 189 | (43) | 177 | (41) |
| Female | 247 | (57) | 259 | (59) |
| Race and ethnicity, n (%) | | | | |
| White, Hispanic/Latino | 2 | (0.5) | 3 | (0.7) |
| White, not Hispanic/Latino | 407 | (93) | 403 | (92) |
| Black, Hispanic/Latino | 1 | (0.2) | 0 | (0) |
| Black, not Hispanic/Latino | 16 | (3.7) | 22 | (5.1) |
| Asian | 1 | (0.2) | 0 | (0) |
| Native American or Alaskan Native | 3 | (0.7) | 1 | (0.2) |
| Multiracial | 1 | (0.2) | 5 | (1.2) |
| [Missing] | 5 | | 2 | |
| Marital status, n (%) | | | | |
| Married or partnered | 292 | (67) | 296 | (68) |
| Divorced or separated | 89 | (20) | 89 | (20) |
| Widowed | 40 | (9.2) | 32 | (7.3) |
| Never married | 14 | (3.2) | 19 | (4.4) |
| [Missing] | 1 | | 0 | |
| Body weight, mean ± SD | 79 | ± 16 | 85 | ± 20 |
| Body weight category, n (%) | | | | |
| <60kg | 45 | (10) | 34 | (7.9) |
| 60–69kg | 90 | (21) | 63 | (15) |
| 70–79kg | 95 | (22) | 89 | (21) |
| 80–89kg | 86 | (20) | 83 | (19) |
| >=90kg | 117 | (27) | 164 | (38) |
| [Missing] | 3 | | 3 | |
| Body mass index, mean ± SD | 26.8 | ± 4.77 | 28.3 | ± 5.47 |
| Body mass index categories, n (%) | | | | |
| Underweight (BMI<18.5) | 5 | (1.2) | 2 | (0.5) |
| Healthy weight (18.5 ≤ BMI < 25.0) | 149 | (35) | 125 | (29) |
| Overweight (25.0 ≤ BMI < 30.0) | 194 | (45) | 167 | (39) |
| Obese I (30.0 ≤ BMI < 35.0) | 60 | (14) | 92 | (21) |
| Obese II (35.0 ≤ BMI < 40.0) | 18 | (4.2) | 31 | (7.2) |
| Obese III (BMI ≥ 40.0) | 5 | (1.2) | 16 | (3.7) |
| [Missing] | 5 | | 3 | |
| Smoking history, n (%) | | | | |
| Current smokers: | | | | |
| 30–42 pack-years | 36 | (8.3) | 77 | (18) |
| 43–53 pack-years | 50 | (11) | 66 | (15) |
| 54–75 pack-years | 45 | (10) | 47 | (11) |
| 76–232 pack-years | 65 | (15) | 51 | (12) |
| Former smokers: | | | | |
| 30–42 pack-years | 40 | (9.2) | 56 | (13) |
| 43–53 pack-years | 59 | (14) | 57 | (13) |
| 54–75 pack-years | 71 | (16) | 45 | (10) |
| 76–232 pack-years | 70 | (16) | 37 | (8.5) |
| VAT measures, median (IQR) | | | | |

| | | | | |
|---|---|---|---|---|
| VAT mean | -88.8 | (-93.2, -84.0) | -89.2 | (-92.6, -85.2) |
| [Missing], n | 2 | | 4 | |
| VAT index | 16.7 | (8.0, 26.7) | 17.4 | (9.6, 29.6) |
| [Missing], n | 3 | | 0 | |
| VAT area | 49.3 | (23.9, 81.0) | 51.1 | (28.9, 91.7) |
| [Missing], n | 0 | | 0 | |
| SAT measures, median (IQR) | | | | |
| SAT mean | -92.6 | (-98.9, -88.0) | -93.3 | (-98.9, -88.0) |
| SAT index | 154.7 | (112.0, 220.0) | 165.5 | (123.4, 230.5) |
| [Missing], n | 6 | | 4 | |
| Muscle measures, median (IQR) | | | | |
| Muscle mean | 28.9 | (24.2, 32.4) | 28.9 | (24.6, 32.4) |
| Muscle index | 137.1 | (117.1, 154.6) | 141.2 | (118.1, 159.8) |
| [Missing], n | 6 | | 4 | |

**Supplemental Table S3.** Primer sequences used for RT-qPCR in mouse muscle and adipose tissues, related to STAR Methods.

| **Mouse Target Gene** | **Forward primer sequence 5'→3'** | **Reverse primer sequence 5'→3'** |
|---|---|---|
| *Fbxo32 (Atrogin 1)* | ATG CAC ACT GGT GCA GAG AG | TGT AAG CAC ACA GGC AGG TC |
| *Atg5* | GAC AAA GAT GTG CTT CGA GAT GTG | GTA GCT CAG ATG CTC GCT CAG |
| *Pnpla2 (Atgl)* | ATG TTC CCG AGG GAG ACC AA | GAG GCT CCG TAG ATG TGA GTG |
| *Becn1* | TTA CCA CAG CCC AGG CGA AA | CTG TAG ACA TCA TCC TGG CTG GG |
| *B2m* | CAT GGC TCG CTC GGT GAC | CAG TTC AGT ATG TTC GGC TTC C |
| *Ccl2 (Mcp1)* | GCT GGA GAG CTA CAA GAG GAT | ACA GAC CTC TCT CTT GAG CTT GGT |
| *CD36* | TCC TCT GAC ATT TGC AGG TCT ATC | AAA GGC ATT GGC TGG AAG AA |
| *Cidea* | AAA GGG ACA GAA ATG GAC AC | TTG AGA CAG CCG AGG AAG |
| *Cxcl1 (KC)* | GCT GGG ATT CAC CTC AAG AA | TGG GGA CAC CTT TTA GCA TC |
| *Gapdh* | ACG ACC CCT TCA TTG ACC TC | TTC ACA CCC ATC ACA AAC AT |
| *Lipe (Hsl)* | TGG CAC ACC ATT TTG ACC TG | TTG CGG TTA GAA GCC ACA TAG |
| *Il6* | CCC GGA GAG GAG ACT TCA CAG | GAG CAT TGG AAA TTG GGG TA |
| *Lep (Leptin)* | TGG CTT TGG TCC TAT CTG TC | TCC TGG TGA CAA TGG TCT TG |
| *Murf1* | TGG CGA TTG TCA CAA AGT GG | CCC TCT CTA GGC CAC CGA GT |
| *Perilipin* | GGG ACC TGT GAG TGC TTC C | GTA TTG AAG AGC CGG GAT CTT TT |
| *Ppia* | GGC AAA TGC TGG ACC AAA C | CAT TCC TGG ACC CAA AAC G |
| *Tnfa* | TCC CAG GTT CTC TTC AAG GGA | GGT GAG GAG CAC GTA GTC GG |
| *Ucp1* | CTC CAG TGG ATG TGG TAA AA | TTC AAA GCA CAC AAA CAT GA |
